# A Path Integral Approach for Allele Frequency Dynamics Under Polygenic Selection

**DOI:** 10.1101/2024.06.14.599114

**Authors:** Nathan W. Anderson, Lloyd Kirk, Joshua G. Schraiber, Aaron P. Ragsdale

## Abstract

Many phenotypic traits have a polygenic genetic basis, making it challenging to learn their genetic architectures and predict individual phenotypes. One promising avenue to resolve the genetic basis of complex traits is through evolve-and-resequence experiments, in which laboratory populations are exposed to some selective pressure and trait-contributing loci are identified by extreme frequency changes over the course of the experiment. However, small laboratory populations will experience substantial random genetic drift, and it is difficult to determine whether selection played a roll in a given allele frequency change. Predicting how much allele frequencies change under drift and selection had remained an open problem well into the 21^st^ century, even those contributing to simple, monogenic traits. Recently, there have been efforts to apply the path integral, a method borrowed from physics, to solve this problem. So far, this approach has been limited to genic selection, and is therefore inadequate to capture the complexity of quantitative, highly polygenic traits that are commonly studied. Here we extend one of these path integral methods, the perturbation approximation, to selection scenarios that are of interest to quantitative genetics. In particular, we derive analytic expressions for the transition probability (i.e., the probability that an allele will change in frequency from *x*, to *y* in time *t*) of an allele contributing to a trait subject to stabilizing selection, as well as that of an allele contributing to a trait rapidly adapting to a new phenotypic optimum. We use these expressions to characterize the use of allele frequency change to test for selection, as well as explore optimal design choices for evolve-and-resequence experiments to uncover the genetic architecture of polygenic traits under selection.

## 1 Introduction

A central problem in evolutionary biology is determining the frequency and genetic basis of adaptive evolution. Two general frameworks within population and quantitative genetics have been offered to study this problem (Stephan, 2016; Höllinger *et al*., 2019). The first proposes that adaptation progresses via a series of substitutions, in which beneficial mutations rise in frequency and fix in the population. Such selective sweeps are expected to distort patterns of local variation, so recent episodes of positive selection can be inferred by scanning the genome for distinctive patterns of diversity (Smith and Haigh, 1974; Hudson *et al*., 1994) or linkage disequilibrium (Kim and Nielsen, 2004) in present-day populations. This approach, founded in population genetics, links an adaptive substitution directly to its effect on fitness, typically omitting consideration of a phenotype under selection except for *post hoc* interpretation.

The second framework is grounded in quantitative genetics, with the understanding that selection acts on phenotypes which affect fitness and that many phenotypic traits have a polygenic basis. When selection occurs on a complex trait, the genetic basis of adaptation may be quite distinct from a classic selective sweep. Instead, the mean phenotype of the population can change rapidly through subtle shifts in frequencies of alleles underlying the trait (Höllinger *et al*., 2019; Barghi *et al*., 2020). While certain genetic architectures, such as low polygenicity and mutations of large effect, can give rise to sweep-like behavior and fixation of trait-affecting alleles (Thornton, 2019; Höllinger *et al*., 2023), power can still be low for identifying signatures of selection. Inferring selection on the alleles underlying a trait under polygenic selection, or even whether polygenic selection has occurred on a given trait in recent history, remains an open and difficult problem in evolutionary genetics (Racimo *et al*., 2018; Novembre and Barton, 2018; Berg *et al*., 2019; Sohail *et al*., 2019).

Time-series data, in which individuals are sampled over multiple generations, has recently emerged as a promising avenue for disentangling the genetic architecture of polygenic adaptation. This includes the rapidly growing body of ancient DNA (Rasmussen *et al*., 2010; Reich *et al*., 2010; Mathieson *et al*., 2015; Mathieson and Terhorst, 2022), the sequencing of museum collections (Card *et al*., 2021; Shpak *et al*., 2023), and evolve and resequence (E&R) experiments (Kawecki *et al*., 2012; Long *et al*., 2015; Schlötterer *et al*., 2015). From these time-series datasets, we gain direct insight into how allele frequencies changed over time and the potential to identify selected alleles from their frequency trajectories. In practice, however, it can be difficult to determine whether a given allele frequency change is the result of selection or solely due to random genetic drift.

Theoretically, the transition density of Kimura’s diffusion equation (Kimura, 1964; Ewens, 2004; Rice, 2004) describes the evolution of allele frequencies under the combined effects of drift and selection. Several existing tests for selection using time-series genomic data appeal to these transition densities (e.g.,Bollback *et al*., 2008; Steinrücken *et al*., 2014; Tataru *et al*., 2015).

Because drift should be random in direction, it is possible to differentiate drift and selection by searching for concordant allele frequency change (AFC) among the replicate populations (Vlachos *et al*., 2019). However, this approach is typically infeasible for natural populations or even smaller experiments (Bull *et al*., 1997; Bollback *et al*., 2008; Barrick *et al*., 2009; Parts *et al*., 2011; Schraiber *et al*., 2016). There has been growing interest in using E&R to study the genetic architecture of phenotypic adaptation. In particular, E&R experiments can be used to assess the degree of parallelism in evolution, that is, whether the same or different alleles respond to selection across replicate populations. For this, selected alleles are identified within each population and the overlap of selected alleles is compared across the full experiment. To identify candidate selected alleles within a population, we may define an AFC threshold based on comparisons to a control population (Parts *et al*., 2011) or neutral simulations (Turner *et al*., 2011; Stern *et al*., 2022) or as an arbitrary value (Bull *et al*., 1997; Barrick *et al*., 2009; Barghi *et al*., 2019; Steyn *et al*., 2023), and then determine the alleles whose frequency changes meet or exceed this threshold.

In E&R experiments, effective population sizes can be small (Keightley and Bulfield, 1993; Johansson *et al*., 2010; Turner *et al*., 2011; Mallard *et al*., 2018; Barghi *et al*., 2019; Claire *et al*., 2021), so that drift will be strong and even neutral alleles can experience extreme AFCs. Insight into how much AFC can be expected under drift, with or without selection, is gained from allelic transition densities. While the transition density of a neutral allele was solved by Kimura (1955b), finding solutions for the transition density under more general selection models has proved challenging. Analytic solutions have only been found in some special cases (Song and Steinrücken, 2012; Steinrücken *et al*., 2013; Schraiber, 2014; Živković *et al*., 2015; Balick, 2023). Most approaches assumed time-invariant selection, though Steinrücken *et al*. (2016) allow for piece-wise constant selection. However, the strength of selection experienced by an allele contributing to a selected polygenic trait does not remain constant as the trait becomes adapted to a new environment. Rather, selection is a function of both time and frequency (Hayward and Sella, 2022).

Here we solve the allelic transition density for two models of polygenic selection. In the first, a population adapts to a new optimum phenotype after a sudden shift in environment (Charlesworth, 2013; Vladar and Barton, 2014; Hayward and Sella, 2022). In the second, a population experiences stabilizing selection around an optimum phenotype in a constant environment (Robertson, 1955; Keightley and Hill, 1988; Simons *et al*., 2018). In this article we focus on the first model, which captures the initial phase of adaptation during which the mean phenotype of a population approaches a new optimum, resulting in time-dependent directional selection on alleles contributing to the trait. We first **(1.1)** review previous work on allelic transition densities, then **(2)** detail the selection model, **(3)** derive our main results, **(4)** explain our simulation and numerical methods, **(5)** show convergence properties, comparing to existing solutions and simulations, **(6)** provide insight on the design of E&R experiments, and finally **(7)** offer concluding remarks.

### 1.1 Background on allelic transition densities

Our main result is solving for allelic transition densities under selection scenarios relevant to selection on a polygenic trait. The transition density,*P*_*α*_(*y, t*|*x*,0), is the probably that a focal allele (here, a trait-contributing allele with a effect size *α*) transitions from frequency *x* at time 0 to frequency *y* at time *t*, measured in units of 2*N*_*e*_ generations. The allelic transition density is the solution to the diffusion equation,

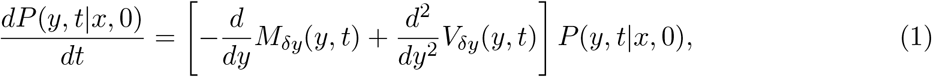

with initial condition *P*(*y*, 0|*x*, 0) = *δ*(*y* − *x*), where *δ* is the Dirac delta function representing an initial known allele frequency of *x*.

The diffusion coefficient *V*_*δy*_(*y, t*) = *y*(1 − *y*)*/*2 describes the contribution to AFC from random genetic drift, and the drift coefficient *M*_*δy*_(*y, t*) = *S*_*α*_(*y, t*) *y*(1 − *y*) describes the contribution to AFC from selection. Other processes such as mutation and migration can also be accounted for in *M*. Here, we only consider selection, and *S*_*α*_(*y, t*) denotes a general selection function that may be time- and frequency-dependent.

Kimura (1955b) solved for the transition probabilities of a neutral allele (denoted *P*_0_(*y, t*|*x*, 0), Equation 2) using the method of eigenfunction expansion. This solution is expressed as an infinite sum of orthogonal functions,

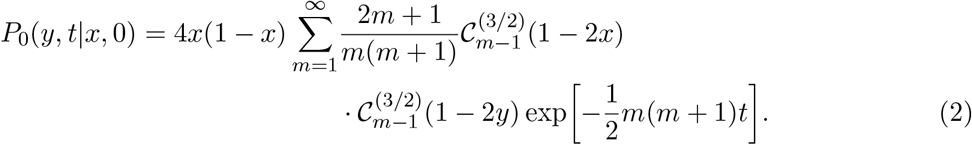

Here, the eigenfunctions are a well known family of polynomials, the Gegenbauer polynomials, denoted 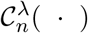. This method was later extended to account for multi-allelic sites and changing demography (Kimura, 1955a) and mutation (Griffiths, 1979).

The transition density of an allele under selection, however, was far more elusive. Kimura (1955c) could not solve for the eigenfunctions exactly, and instead relied on a perturbation expansion of the eigenvalues and functions. This solution is only a viable approximation for weak selection (2*N*_*e*_*s* < 1). Much later, Barbour *et al*. (2000) solved for the selected transition density by expressing allele frequencies as a stochastic process dual to the diffusion. However, this approach is computationally inefficient, bordering on impractical (Steinrücken *et al*., 2013). Song and Steinrücken (2012) reduced the eigenfunctions to an infinite dimensional linear algebra problem, which can be truncated and solved numerically to an arbitrary accuracy even with strong selection (2*N*_*e*_*s* ≫ 1). This numerical method was later extended by Steinrücken *et al*. (2013) to account for multiallelic sites, Živković *et al*. (2015) to account for piece-wise constant demography, and Steinrücken *et al*. (2016) to account piece-wise constant selection.

Schraiber (2014) introduced an alternative approach to solve for the selected transition density. This involves representing the transition probability as a path integral that is then approximated using a perturbation analysis. Recently, Balick (2023) introduced a second path integral method rooted in Langevin equations and solved for transition densities of alleles under strong selection. Our approach follows the path integral method from Schraiber (2014), applied to non-constant selection functions relevant to selection on polygenic traits.

## 2 Selection Model

Our model of the genetic basis of a complex trait assumes that allelic effects combine additively to produce an individual’s phenotype. For highly polygenic traits, this results in a “many-to-one” genotype-phenotype map (Svensson, 2022). Under such a model, two populations could adapt to equivalent environmental changes - a new optimum phenotype - through frequency changes in completely disjoint sets of alleles.

Prior to any change in the phenotypic optimum, we assume that the trait is under stabilizing selection. The strength of stabilizing selection is measured by 1*/W*, where *W* is the width of the fitness function, which we assume to be normal (i.e., Guassian stabilizing selection). Robertson (1955) showed that stabilizing selection results in selection on trait-contributing alleles akin to symmetric underdominance, so that *S*_*α*_(*y, t*) ∝ −(1 − 2*y*)*α*^2^*/*2*W*, that is, selection against the minor allele. This model has historically been very useful for describing the genetic variance and architecture of traits under stabilizing selection (e.g., Keightley and Hill, 1988; Simons *et al*., 2018).

After a sudden shift in optimum phenotype (Figure 1), a focal allele will experience both frequency- and time-dependent selection (Charlesworth, 2013; Vladar and Barton, 2014; Hayward and Sella, 2022). Following Hayward and Sella (2022), for an optimum shift by initial amount Λ, which we assume to be positive, and rescaling time by 2*N*_*e*_,

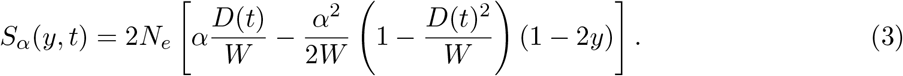

Here, *D*(*t*) is the distance from the population’s mean phenotype to the new optimum (so *D*(0) = Λ), which decreases over time as the population adapts to their new environment. If Λ is large (*O*(*W*)), dynamics are initially dominated by the first term, which behaves as time-variant directional selection. Over time, the strength of directional selection decays and stabilizing selection becomes stronger, so that allelic dynamics once again resemble underdominance.

**Figure 1:**
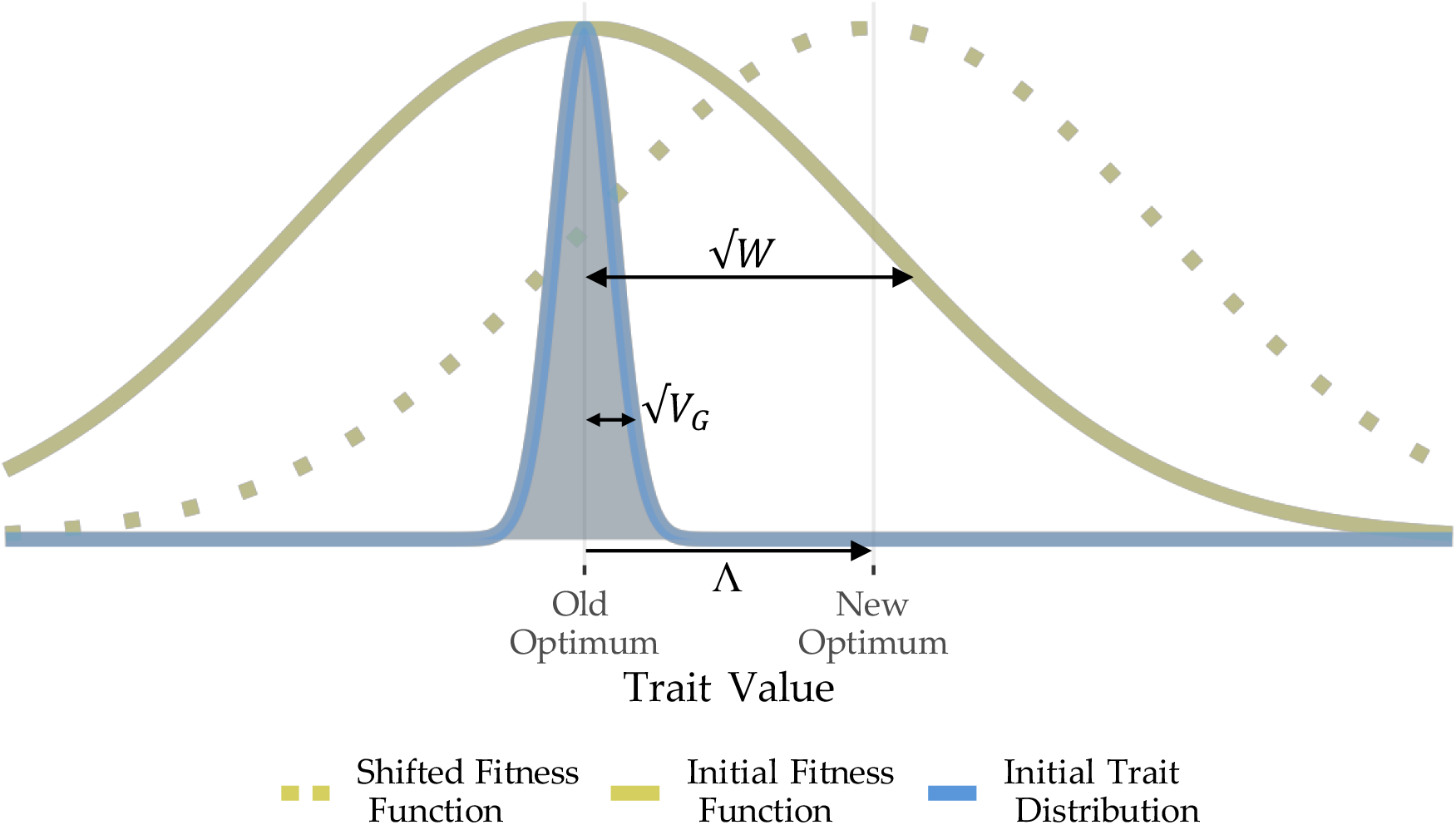
The evolutionary model. Before a shift in optimum, we assume phenotypes are distributed symmetrically around the initial optimal phenotype with variance (*V*_*G*_) that is much smaller than that of the fitness function (*W*). At some time, we consider an instantaneous shift in the optimum trait value (Λ). In the early periods after the optimum shift, dynamics are dominated by directional selection on the trait. This is followed by stabilizing selection to reach a new steady state once the mean phenotype in the population reaches the new optimum (Hayward and Sella, 2022).

For computational efficiency (see Appendix B), we focus on the time-dependent directional selection a focal allele experiences as the trait approaches the new optimum. Thus, we are assuming the trait does *not* reach the new optimum over the time periods considered. Because the time scales of E&R experiments are generally short, this seems a reasonable limitation. In the infinitesimal limit, the approach to the new optimum is well characterized by exponential decay (Lande, 1976; Hayward and Sella, 2022), taking the form *D*(*t*) = Λ exp (−2*N*_*e*_*V*_*G*_*t/W*), where *V*_*G*_ is the additive genetic variance at the time of the optimum shift, and *t* is measured in units of 2*N*_*e*_ generations. Our simplified selection model is then

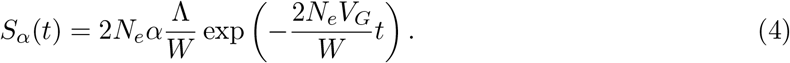

A larger effective population size *N*_*e*_ or allelic effect *α* both lead to stronger selection on the focal allele. In this model, the strength of selection is now a function of the size of the initial change in optimum, Λ, relative to the width of the fitness function, *W*. Larger shifts in optimum phenotype or stronger selection on the phenotype will increase the strength of selection on a focal allele. The rate of decay is controlled by the ratio of *V*_*G*_ to *W*. Either greater genetic variance or a narrower fitness function will increase the rate of adaptation leading to more transient directional selection. We assume that *V*_*G*_ remains constant over the time period modeled here.

In Appendix A.7, we consider the dynamics of focal allele contributing to a trait under constant stabilizing selection. The selection model in this setting is symmetric underdominance, where

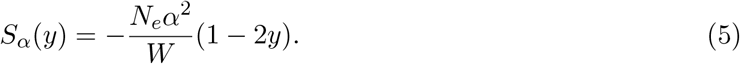

In contrast to the time-dependent selection model immediately after a shift in optimum phenotype (Equation 4), this characterizes the long-term behavior prior to an optimum shift or after the new optimum has been reached. While we do not do so here, it is possible to approximate the full selection model (Equation 3) by dividing time into two phases (Hayward and Sella, 2022). In the first “rapid phase”, the population is adapting to its new environment, and trait contributing alleles experience time-dependent directional selection (Equation 4). In the second “equilibrium phase”, the population has reached its new optimum phenotype and trait contributing alleles experience symmetric underdominant selection (Equation 5).

## 3 Path Integral Solution

A path integral expresses a transition probability, *P*(*y, t*|*x*, 0), by computing the probability of a given allele frequency trajectory (or path) through time, 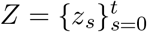, between the initial, *z*_0_ = *x*, and final, *z*_*t*_ = *y*, frequencies, and then integrating over every possible path (Appendix A.1),

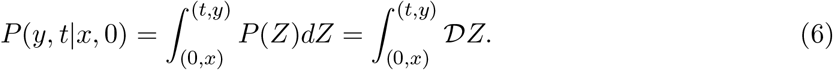

Few path integrals yield exact solutions (Feynman, 1972, Chapter 3.2); however, we can conveniently approximate them using a perturbation scheme. Perturbation analyses are well known in the mathematics literature (Langouche *et al*., 1982; Schulman, 1996; Dickman and Vidigal, 2003) and have been widely used in quantum mechanics (Feynman, 1972, 2010), but are underused in population genetics (but see Rouhani and Barton, 1987; Schraiber, 2014; Balick, 2023). While transition densities may be derived without the use of path integration, we find this formulation particularly intuitive. Here, we outline the steps in deriving our main results, and the full derivation is provided in Appendix A.

As in Schraiber (2014), we begin the perturbation scheme by solving for the ratio of probabilities of a path for a trait-contributing and neutral allele and integrating over all neutral paths

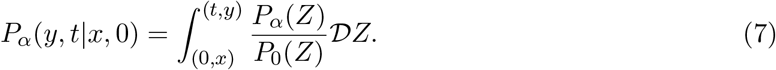

Here, the measure on the path space 𝒟*Z* is induced by the *neutral* wright fisher process. We manipulate the right hand side into the form

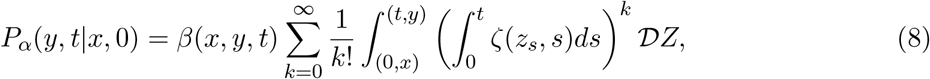

where *β* and *ζ* are functions of the drift and diffusion coefficients, *M* and *V*(Appendix A.2). *β*, which is function of the endpoints *x* and *y* but independent of the path taken between them, captures the deterministic effect of selection. Using terminology from physics, the path integral

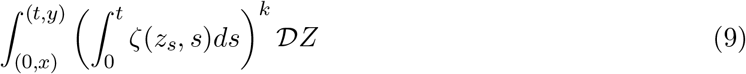

can be interpreted as an allele drifting freely through a selection potential, *ζ*. In other words, the allele neutrally drifts but is scattered by the potential *k* times along its path (Feynman, 1972, Chapter 3.3; Feynman, 2010, Chapter 6-1; Schraiber, 2014). We then integrate over all frequencies and times, *z*_*i*_ and *s*_*i*_, these interactions could take place. For example, with *k* = 1

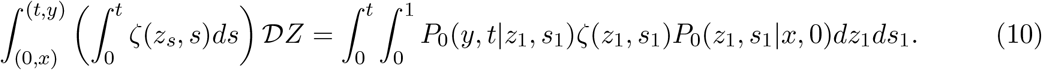

Reading from right to left, this is the probability an allele drifts from its starting frequency *x*, to some arbitrary frequency *z*_1_ at time *s*_1_, is scattered by the potential, then continues to drift to its final frequency *y* at time *t*(Appendix A.3). In general, a *k*^*th*^ order perturbation will lead to products 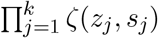 and 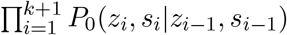.

Piecing these together (Appendix A.4), we find the transition density of a trait contributing allele immediately after a shift in optimum phenotype,

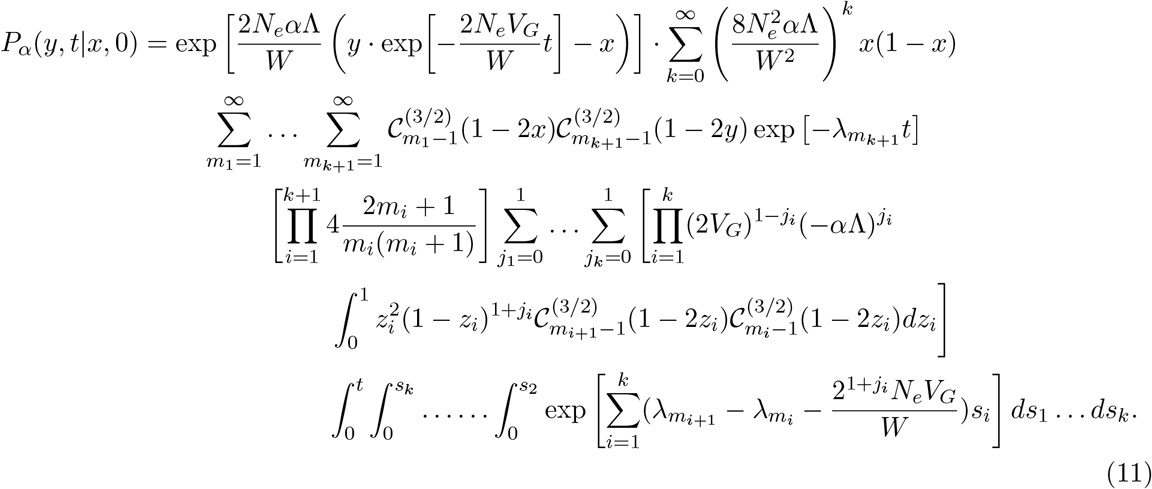

Here, *λ*_*m*_ = *m*(*m* +1)*/*2. The exponent on the first line captures the deterministic effect of selection (*β*, as described above), and the sum over *k* represents the number of times the allele interacts with the selection potential. The remaining terms were introduced by the perturbation analysis. The neutral transition density is an infinite sum (Equation 2) and the series of sums over *m*_1_ … *m*_*k*+1_ result from the product of the *k* + 1 neutral transition densities. Similarly, the selection potential for a trait contributing allele is also a sum of two terms and the series of sums over *j*_1_ … *j*_*k*_ come from the product of *k* selection potentials. Importantly, the integrals over frequency (*z*_*i*_) and time (*s*_*i*_) are separated, and we show in Appendix A.5 and A.6 that these integrals are solvable. In practice, the infinite sums over *k* and *m*_*i*_ are truncated at values of *k*_max_ and *m*_max_, both of which affect the convergence of the solution (Section 5.1).

We find a similar expression for an allele contributing to a trait under constant stabilizing selection, with dynamics described by symmetric underdominant selection on the allele (Equation A44).

An underdominant component appears in the full selection model (Equation 3), and this solution is applicable when there has been no shift in optimum phenotype or for large enough *t* so that the mean phenotype of the population has reached the new optimum. The transition probability for the full selection model (Equation 3) is far more complicated than either of the two simplified models (Appendix B). Indeed, it is more computationally efficient to evaluate *both* simplified models than to evaluate the full model. For this reason, we only fully derive expressions for the two simplified models. In the sections that follow, we focus on the time-dependent directional selection experienced by a focal trait contributing allele immediately after a shift in optimum phenotype.

## 4 Methods

### 4.1 Numerical Methods

To validate our analytic solution for allelic transition density (Equation 11), we numerically solved the diffusion equation (Equation 1) subject to the natural boundary conditions (*sensu* Feller, 1951; c.f. Ewens, 2004, page 149) at 0 and 1 using the NDSolve function in Mathematica 13.2 (Wolfram Research, 2023). We approximated the initial condition, *P*(*y*, 0|*x*, 0) = *δ*(*y* − *x*) with a triangular distribution as implemented in Mathematica 13.2 (Wolfram Research, 2023)

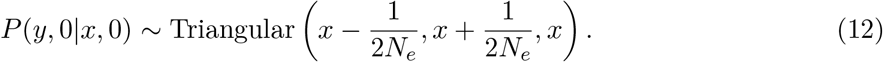

### 4.2 Simulation Methods

In addition to comparisons to the numerical solution (above), we simulated a polygenic trait adapting to a new optimum phenotype as additional validation to our path integral solution. We used two forward-in-time simulation frameworks, one using individual-based simulation of a large chromosome with linkage, and a second assuming linkage equilibrium between all trait-affecting alleles.

For simulations with linkage, we used `fwdpy11` (Thornton, 2014, 2019) to simulate a 1 Morgan genomic region along which mutations appear at a rate *U* mutations per haplotype per generation. Mutation effect sizes were ±*α* with equal probability. While our analytic solution works equally well for negatively selected alleles, we only show results for positively selected alleles (those whose effect sign matches the direction of the optimum shift). We allowed a source population of *N* = 10, 000 individuals to evolve around the initial optimum phenotype for 10*N* generations, at which point allele frequencies were recorded in the source population immediately prior to the establishment of smaller replicate populations. We also calculated linkage disequilibrium among alleles with effects of the same and opposite signs in this generation using `tskit` (Ralph *et al*., 2020). We split each source population into replicate populations of 500 individuals and, after 50 generations of selection, we recorded final allele frequencies in each. We repeated this process 100 times for each parameter combination explored.

Because genetic variance oscillates around its predicted steady state value and classical predictions break down with linkage, we averaged the genetic variance over the last 1,000 generations of the burn-in across independent simulations, which we tracked in `fwdpy11`. We used this value of *V*_*G*_ as the parameter in the analytic solution when directly comparing to simulations. We used the same value for the census size of the simulated populations, *N*, and the effective population size in our analytic solution, *N*_*e*_. Effective sizes of the simulated populations are depressed by selection, but this effect appears to be negligible (Section 5.3). To increase the number of observations in each simulation, we binned together alleles with starting frequencies within ±1% of the input starting frequency in the analytic solution (e.g., we compared simulated alleles with starting frequency 0.09 < *x* < 0.11 to our analytic solution with *x* = 0.1).

We assessed the amount of linked selection experienced by neutral alleles within the 1 Morgan region by adding mutations to the simulated tree sequences using `msprime` (Baumdicker *et al*., 2021). We then compare their AFCs to a completely neutral simulation of the same size using `msprime`, as well as to Kimura’s neutral solution (Equation 2, Kimura, 1955b).

The second set of simulations assumed a completely unlinked genetic architecture. In this simulation, each generation consists of selection under a Gaussian fitness function followed by reproduction and mutation. During the selection phase, each allele independently and deterministically changes in frequency according to

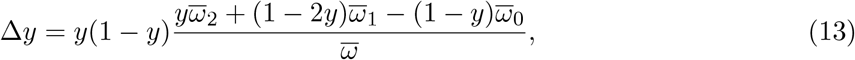

where 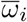 is the marginal fitness of individuals carrying *i* copies of the focal allele and 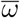 is the mean fitness of the population (Crow and Kimura, 1970, Chapter 5.2), assuming random mating and linkage equilibrium. During the reproduction phase, the next generation’s allele frequency, *y* ^*′*^, is drawn from a binomial distribution

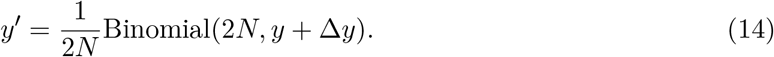

All simulations were performed under an infinite sites model.

### 4.3 Allele Frequency Change (AFC) Threshold Test for Selection

To detect selection, it is common practice to look for alleles with a high amount of allele frequency differentiation between populations in different environments (Turner *et al*., 2010, 2011; Turner and Miller, 2012) or within populations over time (Bull *et al*., 1997; Barrick *et al*., 2009; Parts *et al*., 2011; Barghi *et al*., 2019; Stern *et al*., 2022; Steyn *et al*., 2023). One approach defines the AFC threshold between putatively neutral and selected alleles based on the amount of neutral drift expected given the effective size of the selected populations and the time elapsed (Stern *et al*., 2022). Following this approach, we defined our threshold based on the upper amount of AFC expected for a neutral allele in the selected population.

We solved for the neutral AFC threshold, Ψ_*x*_, such that a neutral allele exceeds that threshold with probability *q*,

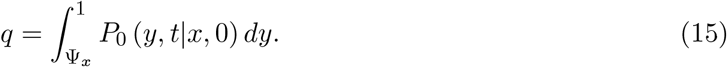

For results shown here, we used *q* = 0.01. The subscript *x* emphasizes the AFC threshold is a function of the starting frequency, but this notation will be neglected later. The specificity of the AFC threshold is 1 − *q* = 0.99. The sensitivity of the AFC threshold is the probability that a selected allele exceeds this threshold and is thus identified as a candidate target of selection,

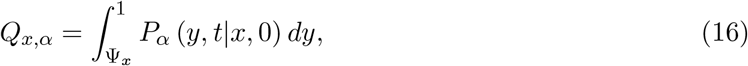

where *P*_*α*_ (*y, t*|*x*, 0) is given by Equation 11. Again, the subscripts are included to emphasize the sensitivity’s dependence on the focal alleles starting frequency and effect (Section 6). The transition probability does not include the probability mass at *y* = 0 or *y* = 1. While the probability an allele fixes by time *t* can be found directly from the transition probability or by solving the Kolmogorov backwards equation (Kimura, 1964), we avoid this complication by limiting our analysis to small *x* and *t* and assuming all lost mass is due to extinction. Integration and solving for Ψ are done in Mathematica 13.2 (Wolfram Research, 2023).

### 4.4 The False Discovery Rate

We assume the starting population is at steady state, and further assume a simple architecture in which all trait-contributing mutations have effect *α* and arise at rate *U*_*α*_ per haploid per generation. Neutral mutations occur at rate *U*_0_. We then show in Appendix C the expected false discovery rate (FDR) among the alleles starting at frequency *x* is

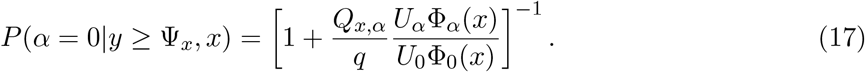

Here, Φ is the density of the mean time a mutation spends at frequency *x* before fixation or loss, which can be found by integrating the transition probability from *t* = 0 to ∞ or by evaluating a general solution (Ewens, 1963; Kimura, 1969; Maruyama, 1974; Ewens, 2004, Chapter 4.4). In particular, the sojourn times of a neutral allele and an underdominant allele are well known (Ewens, 1963, Equation 2.15; Ewens, 2004, Equation 5.18; Simons *et al*., 2018, Equation A18; Yair and Coop, 2022, Equation A12; Hayward and Sella, 2022, Equation S2.2). In Appendix C, we show how to extend this result to an arbitrary distribution of effect sizes and genome-wide scans.

To bound the expected FDR below some value *γ*, we must have

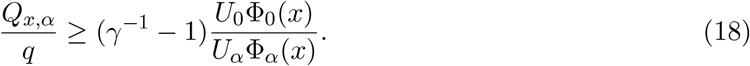

For example, if we want an FDR ≤ 0.05 then *γ*^−1^ − 1 = 19. Furthermore, we expect the ratio on the right-hand side to be ≫ 1 because the target size of the trait under selection is likely much smaller than the size of the genome (*U*_*α*_ ≪ *U*_0_) and Φ_*α*_ < Φ_0_ for nearly all *x*. Thus, the ratio *Q*_*x,α*_*/q* must be very large in order to bound the false discovery rate. As we see later (Section 6; Figure 4), this condition is not typically met. Even when the genetic architecture of the trait consists of few, common loci of large effect, Vlachos *et al*. (2019) found this condition is not met even with the benefit of replicate information. Overall, we should expect a large proportion of the alleles identified as underlying a polygenic trait under selection based on their AFCs to be false positives.

### 4.5 Data availability

Scripts to run simulations, create figures, and evaluate the path integral solution are available at https://github.com/NW-Anderson/transition_probs.

## 5 Results and Discussion

### 5.1 Convergence

There are two series of infinite sums in our solution (Equation 11), representing the number of interactions with the selection potential (Equation 9) and the number of terms from Kimura’s neutral solution (Equation 2), respectively. In practice, these sums are truncated at values *k*_max_ and *m*_max_. Convergence of our solution depends on both values. For stronger selection, smaller genetic variance, and/or longer times, greater *k*_max_ is needed (Figure S1–S4). Convergence for long times depends very little on the value of *m*_max_ (Figure S2), while convergence for small times is dominated by *m*_max_, with more terms required for greater selection (Figure S3).

We quantify the error of our path integral solution using the “total variation” or “statistical” distance between the path integral solution, 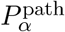, and the numerical solution, 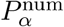, as

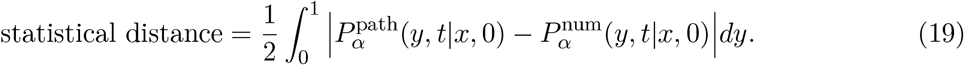

Informally, the statistical distance is half the area bounded between the two distributions.

We find that error is greater under strong selection and low genetic variance, and for longer times (Figures S9, S10). The starting frequency only has an appreciable effect on error when *V*_*G*_ is large. As expected, the error decreases for greater values of *k*_*max*_ (Figure S10). Our solution is unstable for very small times (*t* ≲ 0.02)). This is because our solution relies on Kimura’s neutral solution (Equation 2), which converges slowly for small times (Ewens, 2004, page 162).

### 5.2 Comparison to Existing Solutions

We compared the transition density of an allele contributing to the selected trait to that of a neutral allele, and to an allele under constant genic selection with the same initial strength of selection (2*N*_*e*_*α*Λ*/W* = 2*N*_*e*_*s* = 5). Figure 2 shows the polygenic selection model approaches the behavior of the constant genic selection model (Schraiber, 2014) when *V*_*G*_ is small, and it is well approximated by the neutral model (Kimura, 1955b) when *V*_*G*_ is large. This is expected as the rate of decay in *S*_*α*_(*t*) (Equation 4) depends on the existing genetic variance. When *V*_*G*_ is large, the population approaches the new phenotypic optimum more quickly, and the directional selection felt by a given trait contributing allele is more transient. On the other hand, when *V*_*G*_ is small, the strength of selection decreases very little over the time elapsed and is well approximated by the constant genic model. Intermediate *V*_*G*_, which is likely more common for most polygenic traits, result in behavior intermediate between these two extremes.

**Figure 2:**
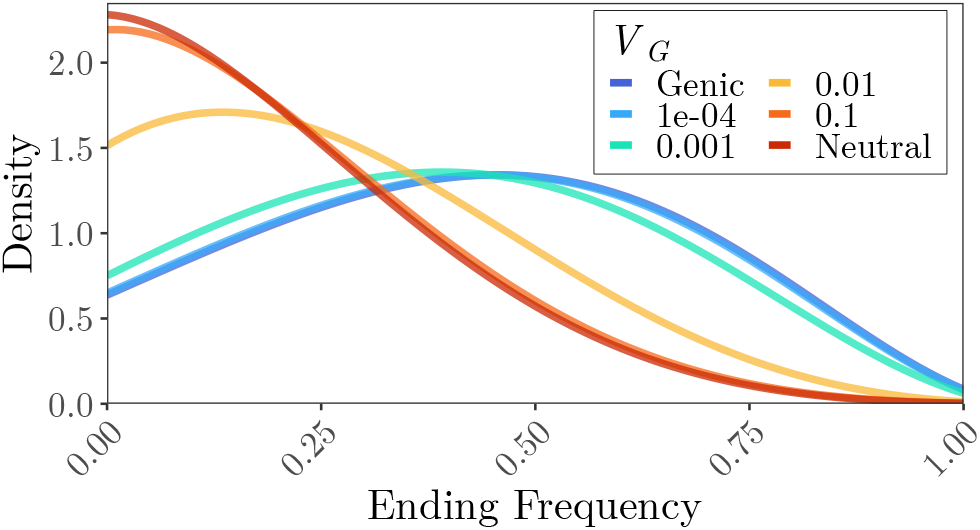
Comparison to existing solutions. The probability distribution of the final frequency of a trait contributing allele with initial frequency *x* = 0.2 and *α* = 0.005 (initial strength of selection 2*N*_*e*_*α*Λ*/W* = 5). Different amounts of genetic variance, which controls the rate of decay in selection, is shown in different colors. Also shown is the neutral solution (Kimura, 1955b) and the genic solution (Schraiber, 2014) with (2*N*_*e*_*s* = 5). When *V*_*G*_ is small (≈10^−4^), the strength of selection decays very slowly, and the allelic dynamics are well approximated by the constant genic solution. At the other extreme, when *V*_*G*_ is large (≈ 10^−1^) selection decays very quickly and the behavior of a focal trait contributing allele is approximately neutral. Solutions were found with parameters *α* = 0.005, *N*_*e*_ = 500, *t* = 0.25, *W* = 1, Λ = 1, *k*_max_ = 4, *m*_max_ = 50.

### 5.3 Comparison to Simulations

Our solution matches the allele frequency dynamics found through simulation of polygenic selection far better than existing solutions which assume genic selection. However, compared to individual-based simulations with linkage, our solution overestimates the AFC of a focal positively selected allele, due to linkage disequilibrium with other selected alleles (Figure 3A). This violates the assumption that the focal allele is in linkage equilibrium with the genomic background. In our simulations with linkage, we observe that alleles of opposite sign tend to be in positive linkage disequilibrium before the shift in optimum (Bulmer, 1971; Figure 3C). In contrast, alleles with same-signed effects tend to be in negative LD with each other. This effect is not present in our simulations without linkage, and we observe excellent agreement with our analytic solution (Figure 3B).

**Figure 3:**
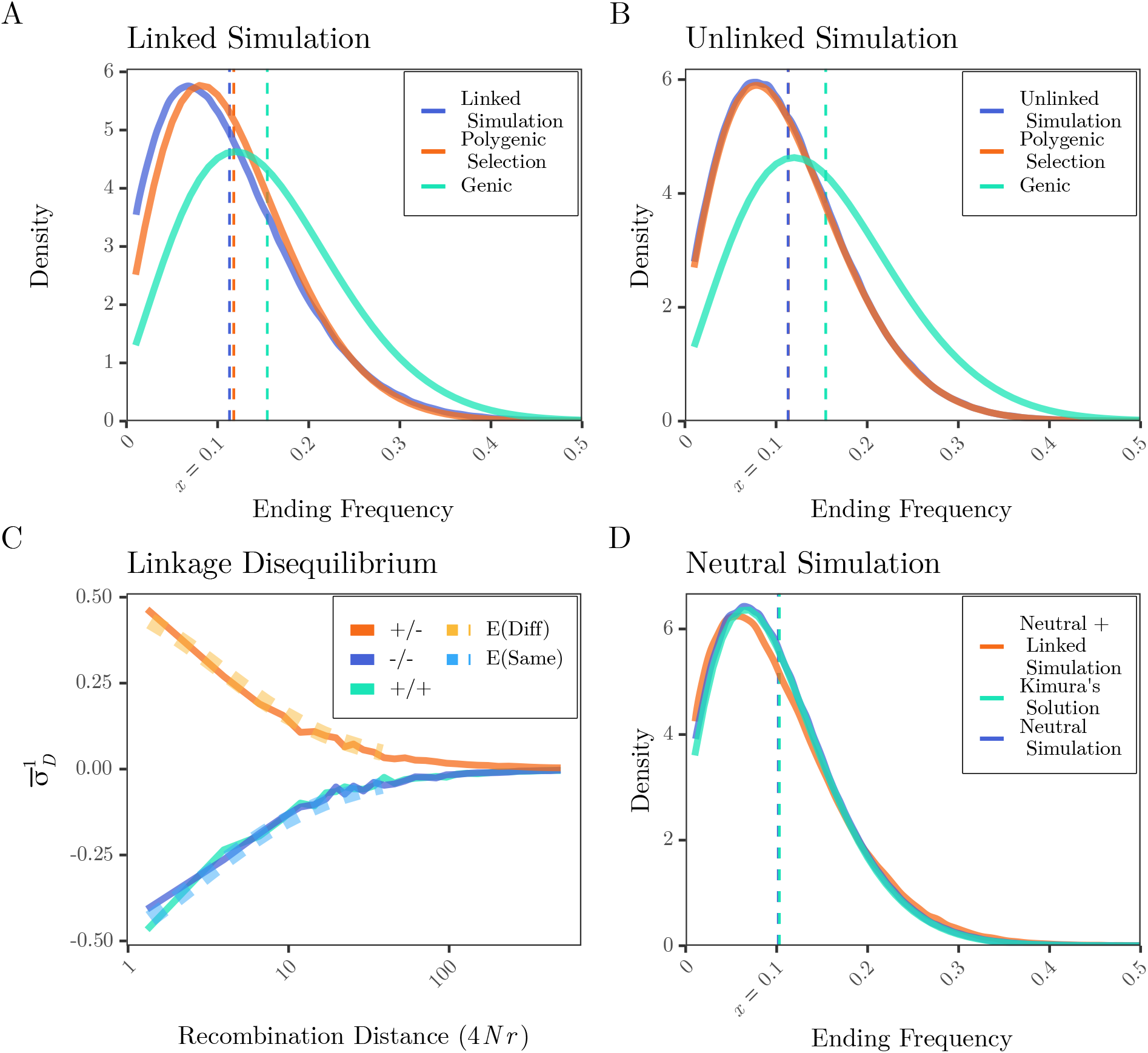
Our solution approximates the allele frequency dynamics found in both linked an unlinked simulations of polygenic selection. (A,B) Distributions of ending frequencies of a trait-contributing allele with initial frequency *x* = 0.1 and *α* = 0.01 (2*N*_*e*_*α*Λ*/W* = 10), after 50 generations (*t* = 0.05) of selection. Vertical dotted lines show the expected or empirical mean ending frequencies from the analytic solution or simulations. Also shown, the constant genic solution (Schraiber, 2014) with (2*N*_*e*_*s* = 10) assumes the strength of selection is constant through time, overestimating the AFC. Linkage increases the probability that a focal positively selected allele decreases in frequency due to LD with negatively selected genomic backgrounds and increases the probability that a neutral allele experiences extreme AFC due to linked selection (D). (C) We computed linkage disequilibrium (signed LD, as 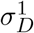 ; Ragsdale, 2022) among alleles both with positive effects (+*/*+), negative effects (−*/*−) and between alleles of opposite effects (+*/*−) as a function of recombination distance measured in the population scaled recombination distance (*ρ* = 4*Nr*) averaged across replicate simulations. Also shown in the dashed lines is the expected LD between alleles of the same and different sign calculated with `moments` (Jouganous *et al*., 2017; Ragsdale, 2022). Model parameters: 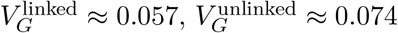, *α* = 0.01, *U* = 0.025, *N* = *N*_*e*_ = 500, *W* = 1, Λ = 1, *k*_max_ = 5, *m*_max_ = 50.

Linkage disequilibrium between trait-affecting alleles reduced the amount of genetic variance by ≈ 20% compared to the unlinked simulations. All else being equal, this decreases the rate the population adapts to the new optimum phenotype and increases the probability of extreme AFCs exceeding a neutral threshold (Figure 4, Section 6.1). Linked selection also has a noticeable effect on neutral allele dynamics. The distribution of ending frequencies of neutral alleles in the selected 1 Morgan region is overdispersed compared to predictions from Kimura’s neutral solution (Kimura, 1955b) and neutral simulations (Figure 3D). The effects of linkage is greatest for higher mutation rates and effect sizes (Figures S5–S7).

**Figure 4:**
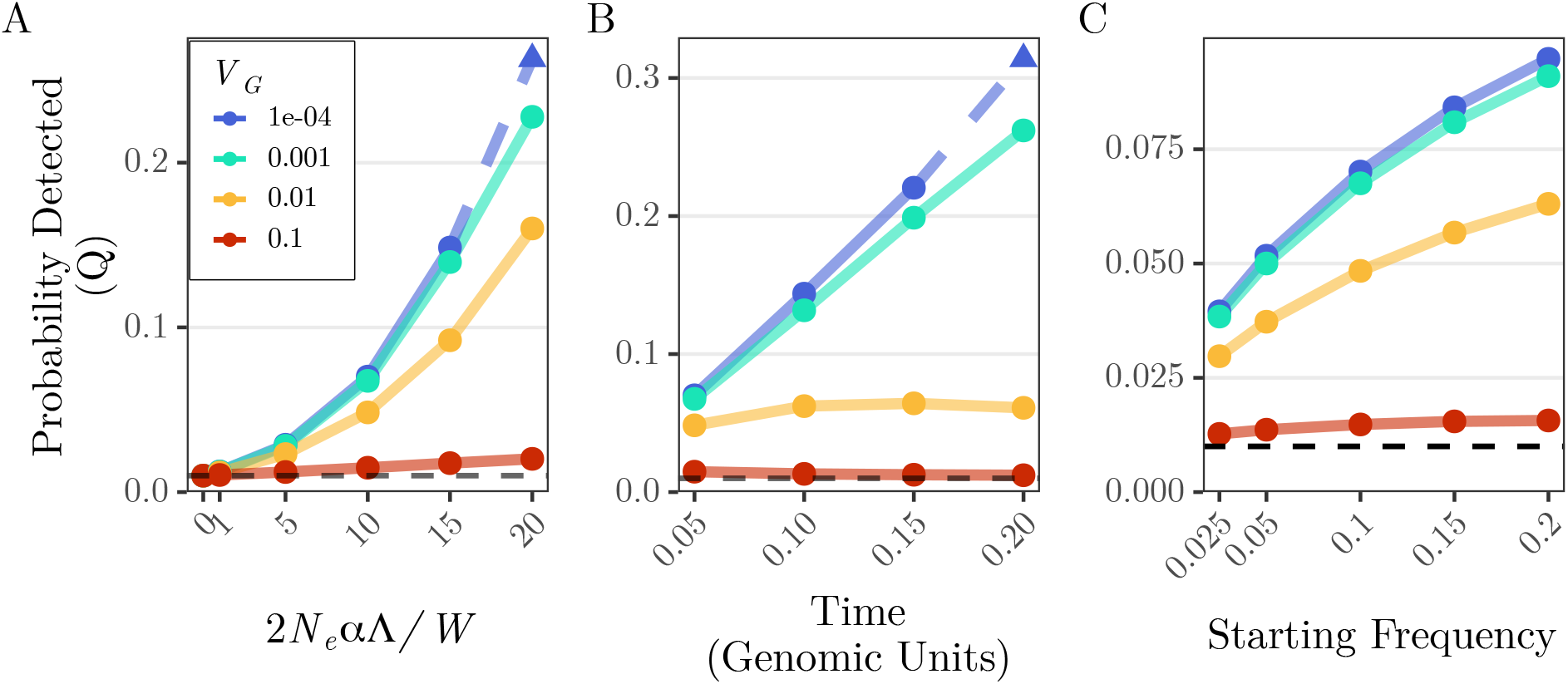
Probability that an allele with 2*N*_*e*_*α*Λ*/W* = 10 exceeds a 99% neutral AFC threshold after 50 generations (*t* = 0.05) with starting frequency *x* = 0.1, unless otherwise stated. (A) The initial strength of selection, (B) the time elapsed, and (C) the starting frequency are varied along the x-axis. The probability is greatest for strong selection, high starting frequency, low genetic variance, and an intermediate time that depends on *V*_*G*_. The horizontal dotted line is the probability that a neutral allele will exceed the AFC threshold, and be falsely flagged as under selection (*q* = 0.01). Triangles and dotted lines indicate numerical results were used due to high error (see Supplemental Figure S9). Model parameters: *N*_*e*_ = 500, *W* = 1, Λ = 1, *k*_max_ = 5, *m*_max_ = 50.

## 6 Allele Frequency Change Threshold Test for Selection

While several studies have explored optimal design choices for E& R experiments, most assume that alleles are under constant genic selection (Kim and Stephan, 1999; Baldwin-Brown *et al*., 2014; Kofler and Schlötterer, 2014). Notable exceptions are Taus *et al*. (2017), who explored the effects of dominance, and Kessner and Novembre (2015), who modeled truncating selection on a quantitative trait. To our knowledge, no studies have explored these design considerations for a polygenic trait under stabilizing selection after a sudden environmental change, despite this being a common experimental design (Teotónio *et al*., 2009; Burke *et al*., 2010; Orozco-terWengel *et al*., 2012; Barghi *et al*., 2019).

### 6.1 The Effect of Genetic Variance

Our polygenic selection model builds upon the genic model by including the decay in the strength of directional selection as the population adapts to their new environment. The rate of decay is controlled by the ratio of genetic variance (*V*_*G*_) and the width of the fitness function (*W*). While, in practice,*W* is an uncontrollable and often unknown parameter,*V*_*G*_ can be inferred (Munar-Delgado *et al*., 2023) and even controlled in the laboratory through inbreeding and outcrossing. We find the probability an allele exceeds our neutral threshold increases with decreasing *V*_*G*_ (Figure 4). In fact, the probability of detecting an allele affecting a trait with high genetic variance is not much greater than the probability a neutral allele exceeds our threshold (Figure 4, red line vs dotted line). With this in mind, if the goal is to detect alleles responding to selection on a quantitative trait with high genetic variance, it may be advisable to inbreed experimental populations to reduce genetic variation prior to the start of the experiment. While some adaptive alleles may be lost, those that remain are more readily detectable. On the other hand, inbreeding can increase linkage disequilibrium which confounds the localization of adaptive variants (Tobler *et al*., 2014; Kofler and Schlötterer, 2014). Furthermore, genetically homogeneous populations can, in reality, go extinct rather than adapt to a new environment. Taking these different factors into account, “intermediate” genetic variances may be the best choice for the identification of the genomic basis of quantitative trait adaptation.

### 6.2 The Effect of Time

Previous studies assuming genic selection have concluded that selected alleles are more easily detected with longer experiments (Kim and Stephan, 1999; Kofler and Schlötterer, 2014).

However, in the polygenic selection model considered here, the strength of selection decreases through time. We find that the probability of detecting a selected allele decreases for longer times when genetic variance is high (Figure 4B). We note that we have assumed the population does not reach its new optimum, which becomes less reasonable for longer periods of time. Once the population reaches the new optimum, stabilizing selection begins to act and trait-contributing alleles with frequency <1*/*2 will experience negative selection, although this selection will tend to be weaker with *s* ∝ *α* ^2^ instead of *α*(Equations 3, 4, 5; Hayward and Sella, 2022). This combination of weaker and and possibly negative selection acting on trait contributing alleles after the population adapts to its new environment would further decrease the probability of detecting a selected allele for long times. For traits with high genetic variance, short experiments with very strong selection would maximize the probability of detecting a selected allele and continuing the experiment after the population has adapted to their new environment would be counterproductive. This result is qualitatively similar to findings by Baldwin-Brown *et al*. (2014); Kessner and Novembre (2015); Taus *et al*. (2017), who found reduced power to detect an allele under constant selection if the experiment is continued after the allele approaches fixation.

### 6.3 The Effect of Starting Frequency

Starting frequency has two conflicting influences on the probability of detection. First, selection does not act efficiently on alleles at frequencies near 0 or 1 (Nei, 1977; Maruyama, 1977, Chapter 4.11). Second, the neutral AFC threshold is greatest for intermediate starting frequencies (the strength of drift *z*(1−*z*)*/*2 is maximized at frequency *z* = 1*/*2). We find alleles starting from high frequency are more probable to exceed the neutral AFC threshold (Figure 4C). This result agrees with Taus *et al*. (2017) under the parameter ranges tested here.

### 6.4 The Effect of Population Size and Number of Replicates

Larger population sizes increase the effective strength of selection relative to random drift. Thus, as observed previously (Kim and Stephan, 1999; Kofler and Schlötterer, 2014; Baldwin-Brown *et al*., 2014; Kessner and Novembre, 2015; Taus *et al*., 2017), the probability of detecting a selected allele increases with increasing population size (Figure S11). So far, we have focused on the probability that an allele will exceed a neutral AFC threshold within a single replicate population. In practice, the genomic parallelism or heterogeneity of adaptation is quantified by how many times each allele is detected among the many replicate populations - the replicate frequency spectrum (RFS; Barghi *et al*., 2019; Stern *et al*., 2022; Steyn *et al*., 2023). We find both the number of replicates and their sizes have large impact on the expected RFS of a large effect allele (*α* = 0.01, 2*N*_*e*_*α*Λ*/W* = 10 when *N*_*e*_ = 500; Figure S12). When populations sizes are small, a trait-contributing allele will exceed the neutral AFC threshold one or only a few times, regardless of the number of replicates. Furthermore, when *N*_*e*_ = 200 the probability of detecting an allele at all exceeds 50% only when the number of replicates exceeds 10. This means, on average, half of large effect (*α* = 0.01) trait contributing alleles will not experience extreme AFCs (relative to the amount of drift) in any of the replicate populations. Alternatively, when the populations are large (*N*_*e*_ = 1000), the probability of detecting an allele with *α* = 0.01 at least once approaches 75% with only 5 replicates, and the mode of the RFS increases with each increase in the number of replicates.

When resources are limited, researchers may be forced to choose between fewer larger replicate populations or a greater number of smaller populations. To explore this situation, we held the total population size in the entire experiment constant, at 5000 individuals, and varied the number of replicates this source population is subdivided into. Thus, doubling the number of replicates halves the size of each one. We find that in this contrived scenario, the AFC threshold returns the fewest false discoveries when there are fewer, larger replicates. Increasing the number of replicates at the expense of their size has the greatest impact on the probability of detecting a neutral allele and falsely labeling it a target of selection (Figure 5). For example, increasing the number of replicates by a factor of 10 from 5 to 50 (with sizes *N*_*e*_ = 1000 and 100, respectively) increases the probability of detecting a neutral allele at least once by ≈ 8×, but yields only a ≈ 6 × increase in the probability for an allele with *α* = 10 ^−3^ and a mere ≈ 1.1 × increase when *α* = 0.01.

**Figure 5:**
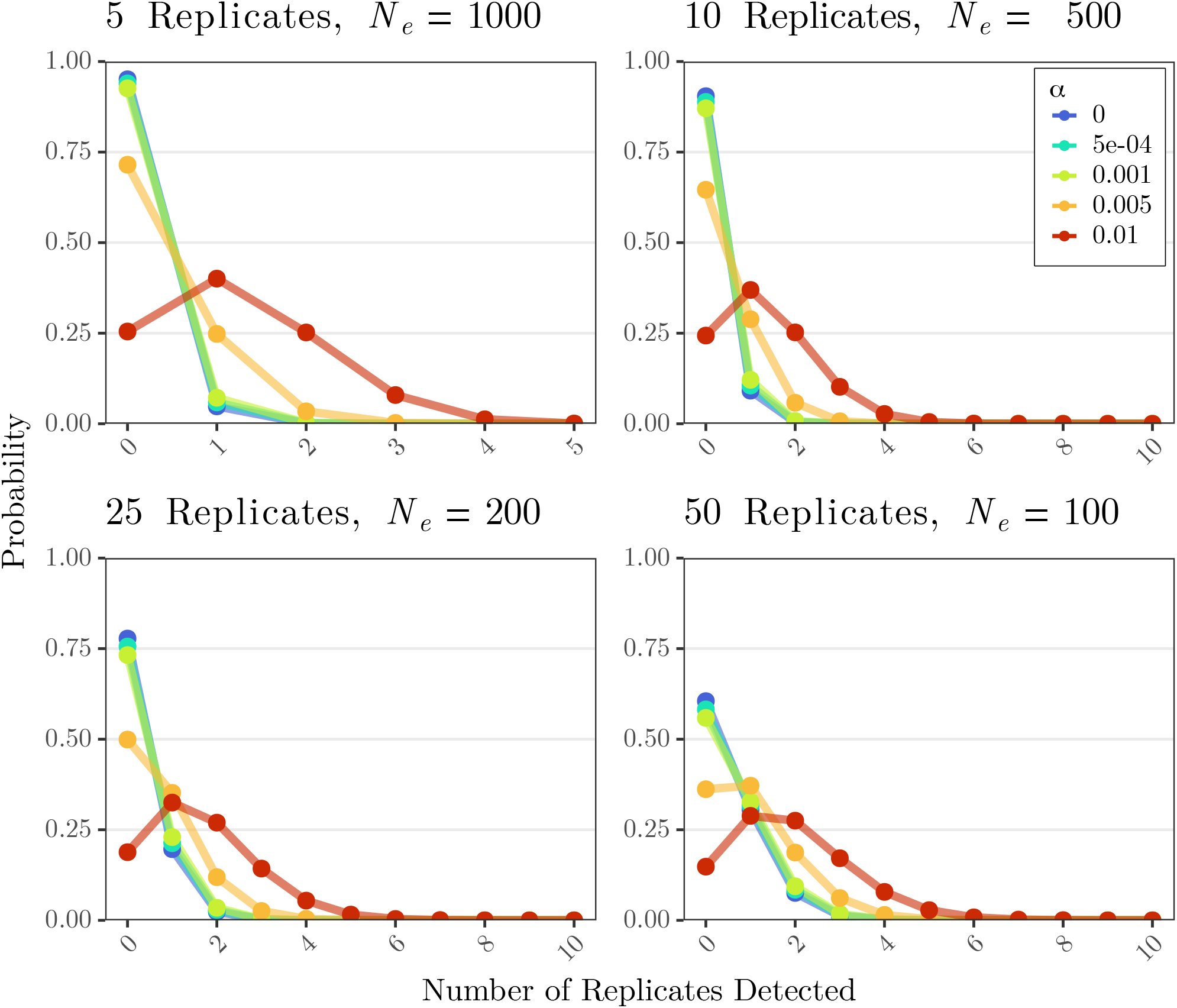
Expected RFS of alleles with effect *α* that exceed a 99% neutral AFC threshold after 50 generations of selection at least once among the replicate populations. The total population size is held constant across the different facets but number and size of the replicate populations is varied. Colors denote the different values of *α* ranging from a neutral to large effect (2*N*_*e*_*α*Λ*/W* = 2 or 20 for *N*_*e*_ = 100 or 1000, respectively). The *x* axis is truncated in the bottom row because the probability of these outcomes is ≈0. Model parameters: *x* = 0.1, *W* = 1, Λ = 1, *V*_*G*_ = 0.001, *k*_max_ = 5, *m*_max_ = 50.

Researchers may adjust the significance threshold, *q*, for multiple comparisons. But, this would also lower the sensitivity of the AFC threshold test for selection. In light of these results, and our previous analysis of the false discovery rate (Section 4.3), we caution against the use of an AFC threshold as a genome wide test for selection especially when replicate information is available. In practice, researchers may first identify candidate alleles under selection using a more powerful test that utilizes replicate information (e.g., the CMH test; Vlachos *et al*., 2019). Then, an AFC threshold is used to determine whether a candidate allele responded to selection within a particular replicate population and to construct an RFS of this subset of candidate alleles (Barghi *et al*., 2019; Stern *et al*., 2022). Because tests that utilize replicate information are typically sensitive to both extreme AFCs in a single replicate as well as smaller AFCs in the same direction among many replicates, the effects of this pre-screening on the RFS are hard to predict and present an interesting problem for future research.

## 7 Conclusions

Path integration continues to be a powerful tool in population genetics. Rouhani and Barton (1987) used path integration to find the rate of transitions in fitness optima in Wright’s “Shifting Balance” theory (Wright, 1931).Mustonen and Lässig (2010) defined the concept of “Fitness Flux” and Neher and Shraiman (2012) derived the rate of Muller’s Ratchet using path integration. Nourmohammad and Eksin (2021) used a path integral control approach to derive artificial selection strategies to guide the evolution of molecular phenotypes. Schraiber (2014) and Balick (2023) derived analytic transition probabilities for alleles under weak and strong genic selection, respectively.

Here, we used a path integral method to derive analytic transition probabilities for an allele contributing to a quantitative trait after a sudden shift in optimum phenotype, resulting in time-dependent selection (Equation 11, A43). We found that this expression well approximates allelic dynamics found through expensive forward-in-time simulation both with and without linkage. This model provided insight into the design of E&R experiments to study the genomic architecture and parallelism of quantitative adaptation. Most notably, we found that longer experiments are less powerful to detect selected alleles contributing to traits with high genetic variance, that lower genetic variance generally increases the probability of detecting a focal trait-contributing allele.

Finally, we used this same path integration approach to find allelic transition densities under constant stabilizing selection, which results in frequency-dependent selection (Equation A44). We leave applications of this expression to future work. While we focus on short term directional selection on a complex trait which is relevant to E&R studies, stabilizing selection is thought to be common (Kingsolver *et al*., 2001; Sanjak *et al*., 2018; Sella and Barton, 2019). Furthermore, stabilizing selection around a fluctuating optimum phenotype has been shown to have important consequences for the maintenance of genetic variation (Ellner and Hairston, 1994; Kawecki, 2000; Bell, 2010; Svardal *et al*., 2011; Villemereuil *et al*., 2020; Johnson *et al*., 2023). The combination of these two settings into a single expression would therefore be a valuable advance for predicting allelic dynamics in these evolutionary scenarios.

## 8 Acknowledgements

We thank Matthias Steinrücken, Gustavo Barosso, and Nick Collier for useful discussion. We also thank Jennifer Jung for valuable comments on the manuscript. NWA was supported through an NHGRI training grant to the Genomic Sciences Training Program 5T32HG002760. Support for this research was provided by the Office of the Vice Chancellor for Research and Graduate Education at the University of Wisconsin–Madison with funding from the Wisconsin Alumni Research Foundation.

## Notes

### Competing Interest Statement

The authors have declared no competing interest.

