## Appendix and Supplemental Figures for "A Path Integral Approach for Allele Frequency Dynamics Under Polygenic Selection"

### A Full Derivation

#### A.1 Introducing the Path Integral

In the next two sections, we will attempt to build some intuition behind path integration and its application to solving transition probabilities. Beginning with some discrete Markov theory, transition probabilities depend only on the current position, with no dependence on past behavior. Transition probabilities are denoted $P(z_s, s|z_{s-1}, s-1)$: the probability that the particle is in state $z_s$ at time $s$, given it was in state $z_{s-1}$ the previous time point. Throughout the paper, $z$ denotes arbitrary allele frequencies ($x$ and $y$ are reserved for the fixed starting and ending frequencies) and $s$ denotes arbitrary times ($t_0$ and $t$ are fixed start and end times, if the start time is not set to 0). The probability that an allele takes a given path $Z = \{z_s\}_{s=0}^{t}$ is the product of the single step probabilities:

$$P(Z) = \prod_{s=1}^{t} P(z_s, s|z_{s-1}, s-1). \tag{A1}$$

Multistep transition probabilities can be calculated by integrating over the possible intermediate steps, known as the Chapman-Kolmogorov Equation:

$$P(z_2, 2|z_0, 0) = \int P(z_2, 2|z_1, 1)P(z_1, 1|z_0, 0)dz_1. \tag{A2}$$

In diffusion processes, there exists infinitely many transitions between any two times. Thus, long time transition probabilities can be calculated as an infinite application of the Chapman-Kolmogorov Equation:

$$\begin{aligned} P(y,t|x,0) = \lim_{N\to\infty} \int \dots \int & P(y,t|z_{N-1}, s_{N-1})P(z_{N-1}, s_{N-1}|z_{N-2}, s_{N-2})dz_{N-1} \\ & \cdot P(z_{N-2}, s_{N-2}|z_{N-3}, s_{N-3})dz_{N-2} \\ & \dots \cdot P(z_1, s_1|x, 0)dz_1 \\ = \int_{(0,x)}^{(t,y)} \mathcal{D}Z, & \end{aligned} \tag{A3}$$

where total time $[0, t]$ is divided into $N$ intervals, with $dt = t/N$, $s_n = ndt$, $z_{s_n} = z_n$ and the endpoints $z_0 = x$ and $z_N = y$. This is the path integral formulation of a transition probability (Feynman, 1972, Chapter 3.2; Risken and Haken, 1989, Chapter 4.4.2; Feynman, 2010, Chapter 2-4). This can be formalized using measure theory, where $\mathcal{D}Z$ is the measure induced by the infinitesimal transition probabilities, and we integrate over every path, $Z$, connecting $(0, x)$ and

$(t, y)$. Using informal notation,

$$P(y,t|x,0) = \lim_{N\to\infty} \int \dots \int P(Z) dz_{N-1}, ..., dz_1 = \int_{(0,x)}^{(t,y)} \mathcal{D}Z \tag{A4}$$

describes the path integral as it integrates over the infinite intermediate steps.

##### A.2 Path Integral Formulation

We solve for the transition density of an allele with selection by solving for the ratio of the probability of a given path with and without selection, then integrating over all neutral paths (Schraiber, 2014)

$$\begin{aligned} P_\alpha(y,t|x,0) &= \int_{(0,x)}^{(t,y)} \lim_{N\to\infty} \frac{\prod_{n=1}^N P_\alpha(z_n, s_n|z_{n-1}, s_{n-1}) dz_n}{\prod_{n=1}^N P_0(z_n, s_n|z_{n-1}, s_{n-1}) dz_n} \mathcal{D}Z \\ &= \int_{(0,x)}^{(t,y)} \rho_{\alpha,0}(Z) \mathcal{D}Z. \end{aligned} \tag{A5}$$

Here, the measure on the path space $\mathcal{D}Z$ is induced by the **neutral** wright fisher process. Intuitively, $\mathcal{D}Z$ gives us the infinite product of neutral transition densities (Equation A3). Then, one can imagine it canceling with the denominator in Equation A5. More formally, this constitutes a change of measure under Girsanov's theorem (Rogers and Williams, 2000, Chapter IV.38) The infinitesimal time transition probabilities of the Diffusion are known in general, the Gaussian Approximation (Risken and Haken, 1989, Chapter 4.4.1),

$$\begin{aligned} P(y, t_0 + ds|x, t_0) &= \frac{1}{\sqrt{2\pi \cdot 2V(x,t_0)ds}} \cdot \exp\left[ -\frac{(y - (x + M(x,t_0)ds))^2}{2 \cdot 2V(x,t_0)ds} \right] \\ &\sim \text{Normal}\left(x + M(x,t_0)ds, 2V(x,t_0)ds\right). \end{aligned} \tag{A6}$$

Substituting this expression into the transition probabilities in Equation A5 we find the ratio

$$\begin{aligned} \rho_{\alpha,0}(Z) &= \lim_{N\to\infty} \frac{\prod_{n=1}^N P_\alpha(z_n, s_n|z_{n-1}, s_{n-1}) dz_n}{\prod_{n=1}^N P_0(z_n, s_n|z_{n-1}, s_{n-1}) dz_n} \\ &= \lim_{N\to\infty} \prod_{n=1}^N \exp\left[ -\frac{[z_n - (z_{n-1} + S_\alpha(z_{n-1} s_{n-1}) z_{n-1}(1 - z_{n-1})ds)]^2 - [z_n - z_{n-1}]^2}{2 z_{n-1}(1 - z_{n-1})ds} \right] \\ &= \lim_{N\to\infty} \exp\left[ \sum_{n=1}^N S_\alpha(z_{n-1}, s_{n-1})(z_n - z_{n-1}) - \frac{1}{2} \sum_{n=1}^N S_\alpha^2(z_{n-1}, s_{n-1}) z_{n-1}(1 - z_{n-1}) ds \right]. \end{aligned} \tag{A7}$$

Let $z_n - z_{n-1} = dz_n$ and take the continuous limit $N \to \infty$

$$\rho_{\alpha,0}(Z) \approx \exp\left[ \int_0^t S_\alpha(z_s, s) dz_s - \frac{1}{2} \int_0^t S_\alpha^2(z_s, s) z_s(1 - z_s) ds \right]. \tag{A8}$$

It is clear from the first sum in Equation A7 that

$$\int_0^t S_\alpha(z_s, s) dz_s \tag{A9}$$

is an Ito stochastic integral. These must be solved using Ito's Formula, as $z_s$ is almost nowhere differentiable (due to the dependence on Brownian motion Equation A11). The integral form of Ito's lemma is

$$\int_0^t \frac{\partial f}{\partial z} dz_s = f(y,t) - f(x,0) - \int_0^t \left( \frac{\partial f}{\partial s} + \frac{\sigma^2}{2} \frac{\partial^2 f}{\partial z^2} \right) ds. \tag{A10}$$

$\sigma^2$ is the stochastic term in SDE

$$dZ_s = \mu(Z_s, s) ds + \sigma(Z_s, s) dB_s, \tag{A11}$$

which is equivalent to the diffusion with

$$M(z,s) = \mu(z,s) \tag{A12}$$

$$2V(z,s) = (\sigma(z,s))^2, \tag{A13}$$

(Gardiner, 2004, Chapter 4.3.5).

Denoting

$$\int_0^t S_\alpha(z_s, s) dz_s = \frac{2N_e \alpha \Lambda}{W} \int_0^t \exp\left[ -\frac{2N_e V_G}{W} s \right] dz_s = \frac{2N_e \alpha \Lambda}{W} \int_0^t \frac{d}{dz} f(z,s) dz_s, \tag{A14}$$

then,

$$f(z,s) = \int^z \frac{d}{dz} f(\xi, s) d\xi = \exp\left[ -\frac{2N_e V_G}{W} s \right] \int^z d\xi = z \cdot \exp\left[ -\frac{2N_e V_G}{W} s \right] \tag{A15}$$

$$\frac{d}{ds} f(z,s) = \left( -\frac{2N_e V_G}{W} \right) \cdot z \cdot \exp\left[ -\frac{2N_e V_G}{W} s \right] \tag{A16}$$

$$\frac{d^2}{dz^2} f(z,s) = 0. \tag{A17}$$

Applying Ito's lemma (Equation A10), we find

$$\int_0^t S_\alpha(z_s, s) dz_s = \frac{2N_e \alpha \Lambda}{W} \left( y \cdot \exp\left[ -\frac{2N_e V_G}{W} t \right] - x + \frac{2N_e V_G}{W} \int_0^t z_s \exp\left[ -\frac{2N_e V_G}{W} s \right] ds \right), \tag{A18}$$

and the ratio (Equation A8)

$$\rho_{\alpha,0}(Z) = \exp\left[\frac{2N_e\alpha\Lambda}{W}\left(y\cdot\exp\Big[-\frac{2N_eV_G}{W}t\Big]-x\right)\right]$$
$$\cdot\exp\left[\frac{2N_e^2\alpha\Lambda}{W^2}\int_0^t 2V_Gz_s\exp\Big[-\frac{2N_eV_G}{W}s\Big]-\alpha\Lambda z_s(1-z_s)\exp\Big[-\frac{4N_eV_G}{W}s\Big]ds\right]. \tag{A19}$$

#### A.3 Perturbation Analysis

Taylor expanding the exponent on the second line around $\alpha = 0$

$$\rho_{\alpha,0}(Z) = \exp\left[\frac{2N_e\alpha\Lambda}{W}\left(y\cdot\exp\Big[-\frac{2N_eV_G}{W}t\Big]-x\right)\right]$$
$$\cdot\sum_{k=0}^{\infty}\frac{1}{k!}\left(\frac{2N_e^2\alpha\Lambda}{W^2}\right)^k\left(\int_0^t\eta(z_s,s)ds\right)^k, \tag{A20}$$

where

$$\eta(z_s,s) = 2V_Gz_s\exp\Big[-\frac{2N_eV_G}{W}s\Big]-\alpha\Lambda z_s(1-z_s)\exp\Big[-\frac{4N_eV_G}{W}s\Big].$$

Recalling that this ratio is inside the path integral formulation of a transition probability (Equation A5)

$$\begin{aligned}P_\alpha(y,t|x,0) &= \int_{(0,x)}^{(t,y)}\rho_{\alpha,0}(z)\mathcal{D}Z\\
&\approx \exp\left[\frac{2N_e\alpha\Lambda}{W}\left(y\cdot\exp\Big[-\frac{2N_eV_G}{W}t\Big]-x\right)\right]\\
&\qquad\cdot\int_{(0,x)}^{(t,y)}\sum_{k=0}^{\infty}\frac{1}{k!}\left(\frac{2N_e^2\alpha\Lambda}{W^2}\right)^k\left(\int_0^t\eta(z_s,s)ds\right)^k\mathcal{D}Z\\
&= \exp\left[\frac{2N_e\alpha\Lambda}{W}\left(y\cdot\exp\Big[-\frac{2N_eV_G}{W}t\Big]-x\right)\right]\\
&\qquad\cdot\sum_{k=0}^{\infty}\frac{1}{k!}\left(\frac{2N_e^2\alpha\Lambda}{W^2}\right)^k\int_{(0,x)}^{(t,y)}\left(\int_0^t\eta(z_s,s)ds\right)^k\mathcal{D}Z.\end{aligned} \tag{A21}$$

The final equality can be proven using Fubini's theorem, which we assume holds.

We approximate the path integral

$$\Xi_k = \int_{(0,x)}^{(t,y)}\left(\int_0^t\eta(z_s,s)ds\right)^k\mathcal{D}Z \tag{A22}$$

using a perturbation approach (Feynman, 1972, 2010; Schraiber, 2014). Using the language of physics, the path integral can be interpreted as an allele drifting freely through a selection potential, $\eta$. In other words, the allele neutrally drifts but is scattered by the potential $k$ times along its path.

We then integrate over all frequencies and times, $z_i$ and $s_i$, these interactions could take place. For example, when $k = 0$, the allele does not interact with the selection potential at all along the path, and, because we are integrating over neutral paths, we recover the neutral transition density

$$\Xi_{k=0} = \int_{(0,x)}^{(t,y)} \mathcal{D}Z = P_0(y,t|x,0). \tag{A23}$$

If $k = 1$ then the allele interacts with the selection potential once during its path, and drifts otherwise. We then integrate over the times $s_1$ and positions $z_1$ this interaction could have occurred

$$\Xi_{k=1} = \int_0^t \int_0^1 P_0(z_1,s_1|x,0)\eta(z_1,s_1)P_0(y,t|z_1,s_1)dz_1 ds_1. \tag{A24}$$

Reading from right to left, this is the probability an allele drifts from its starting frequency $x$, to some arbitrary frequency $z_1$ at time $s_1$, is scattered by the potential, then continues to drift to its final frequency $y$ at time $t$. Letting $z_0 = x$, $z_{k+1} = y$, $s_0 = 0$, and $s_{k+1} = t$ for notational convenience, the general form for $\Xi_k$ becomes obvious

$$\Xi_k = k! \int_0^t \int_0^{s_k} \cdots\cdots \int_0^{s_2} \int_0^1 \underbrace{\cdots\cdots}_{k \text{ times}} \int_0^1 \left[\prod_{i=1}^{k+1} P_0(z_i,s_i|z_{i-1},s_{i-1})\right] \cdot \left[\prod_{j=1}^{k} \eta(z_j,s_j)\right] dz_1 \ldots dz_k ds_1 \ldots ds_k, \tag{A25}$$

where the $k!$ comes from the number of possible orderings of the k interactions.

#### A.4 Separating Time and Frequncy Integrals

Next, we separate the integrals over space ($z_i$) and time ($s_j$). We begin with the product of neutral transition probabilities (Equation 2; Kimura, 1955b)

$$\prod_{i=1}^{k+1} P_0(z_i,s_i|z_{i-1},s_{i-1}) = \prod_{i=1}^{k+1}\left[4z_{i-1}(1-z_{i-1})\sum_{m=1}^{\infty}\frac{2m+1}{m(m+1)}\mathcal{C}_{m-1}^{(3/2)}(1-2z_{i-1}) \cdot \mathcal{C}_{m-1}^{(3/2)}(1-2z_i)\exp\left[-\frac{1}{2}m(m+1)(s_i-s_{i-1})\right]\right], \tag{A26}$$

where $\mathcal{C}_{m-1}^{(3/2)}$ are the Gegenbauer polynomials. We can exchange the order of summation and multiplication using the fact

$$\prod_{m=1}^{M}\sum_{n=1}^{N} x_{m,n} = \sum_{n_1=1}^{N}\cdots\sum_{n_M=1}^{N}\prod_{m=1}^{M} x_{m,n_m}. \tag{A27}$$

Thus,

$$\prod_{i=1}^{k+1} P_0(z_i, s_i | z_{i-1}, s_{i-1}) = \sum_{m_1=1}^{\infty} \cdots \sum_{m_{k+1}=1}^{\infty} \prod_{i=1}^{k+1} \Big[ 4 z_{i-1}(1-z_{i-1}) \frac{2m_i+1}{m_i(m_i+1)} \cdot \mathcal{C}_{m_i-1}^{(3/2)}(1-2z_{i-1}) \mathcal{C}_{m_i-1}^{(3/2)}(1-2z_i) \exp[-\lambda_{m_i}(s_i - s_{i-1})] \Big], \tag{A28}$$

where $\lambda_m = \frac{1}{2}m(m+1)$. Combining this and Equations A25 and A21, we find

$$P_\alpha(y,t|x,0) = \exp\left[\frac{2N_e\alpha\Lambda}{W}\left(y \cdot \exp\Big[-\frac{2N_eV_G}{W}t\Big] - x\right)\right] \cdot \sum_{k=0}^{\infty}\left(\frac{2N_e^2\alpha\Lambda}{W^2}\right)^k \int_0^t \int_0^{s_k} \cdots\cdots \int_0^{s_2} \int_0^1 \underbrace{\cdots\cdots}_{k \text{ times}} \int_0^1 \left[\sum_{m_1=1}^{\infty} \cdots \sum_{m_{k+1}=1}^{\infty} \left[\prod_{i=1}^{k+1} z_{i-1}(1-z_{i-1})\right]\left[\prod_{i=1}^{k+1} 4\frac{2m_i+1}{m_i(m_i+1)}\right] \left[\prod_{i=1}^{k+1} \mathcal{C}_{m_i-1}^{(3/2)}(1-2z_{i-1})\right]\left[\prod_{i=1}^{k+1} \mathcal{C}_{m_i-1}^{(3/2)}(1-2z_i)\right] \left[\prod_{i=1}^{k+1} \exp[-\lambda_{m_i}(s_i - s_{i-1})]\right]\right] \left[\prod_{j=1}^{k} \eta(z_j, s_j)\right] dz_1 \ldots dz_k ds_1 \ldots ds_k. \tag{A29}$$

Pulling the endpoints $z_0 = x$, $z_{k+1} = y$, $s_0 = 0$, and $s_{k+1} = t$ out of the products, and reindexing where appropriate

$$
\begin{aligned}
P_\alpha(y,t|x,0) = \exp\left[\frac{2N_e\alpha\Lambda}{W}\left(y\cdot\exp\Big[-\frac{2N_eV_G}{W}t\Big]-x\right)\right] \\
\cdot\sum_{k=0}^{\infty}\left(\frac{2N_e^2\alpha\Lambda}{W^2}\right)^k \int_0^t\int_0^{s_k}\cdots\cdots\int_0^{s_2}\int_0^1\underbrace{\cdots\cdots}_{k \text{ times}}\int_0^1 \\
\left[\sum_{m_1=1}^{\infty}\cdots\sum_{m_{k+1}=1}^{\infty}\left[x(1-x)\prod_{i=1}^{k}z_i(1-z_i)\right]\left[\prod_{i=1}^{k+1}4\frac{2m_i+1}{m_i(m_i+1)}\right]\right. \\
\left[\mathcal{C}^{(3/2)}_{m_1-1}(1-2x)\prod_{i=1}^{k}\mathcal{C}^{(3/2)}_{m_{i+1}-1}(1-2z_i)\right] \\
\left[\mathcal{C}^{(3/2)}_{m_{k+1}-1}(1-2y)\prod_{i=1}^{k}\mathcal{C}^{(3/2)}_{m_i-1}(1-2z_i)\right] \\
\left.\left[\exp\left[-\lambda_{m_{k+1}}t\right]\cdot\prod_{i=1}^{k}\exp\left[(\lambda_{m_{i+1}}-\lambda_{m_i})s_i\right]\right]\right] \\
\left[\prod_{j=1}^{k}\eta(z_j,s_j)\right]dz_1\ldots dz_k ds_1\ldots ds_k.
\end{aligned}
\tag{A30}
$$

After combining all $k$ fold products

$$
\begin{aligned}
P_\alpha(y,t|x,0) = \exp\left[\frac{2N_e\alpha\Lambda}{W}\left(y\cdot\exp\Big[-\frac{2N_eV_G}{W}t\Big]-x\right)\right] \\
\cdot\sum_{k=0}^{\infty}\left(\frac{2N_e^2\alpha\Lambda}{W^2}\right)^k x(1-x)\int_0^t\int_0^{s_k}\cdots\cdots\int_0^{s_2}\int_0^1\underbrace{\cdots\cdots}_{k \text{ times}}\int_0^1 \\
\sum_{m_1=1}^{\infty}\cdots\sum_{m_{k+1}=1}^{\infty}\mathcal{C}^{(3/2)}_{m_1-1}(1-2x)\mathcal{C}^{(3/2)}_{m_{k+1}-1}(1-2y)\exp\left[-\lambda_{m_{k+1}}t\right] \\
\left[\prod_{i=1}^{k+1}4\frac{2m_i+1}{m_i(m_i+1)}\right]\left[\prod_{i=1}^{k}z_i(1-z_i)\mathcal{C}^{(3/2)}_{m_{i+1}-1}(1-2z_i)\mathcal{C}^{(3/2)}_{m_i-1}(1-2z_i)\right. \\
\left(2V_Gz_i\exp\Big[(\lambda_{m_{i+1}}-\lambda_{m_i}-\frac{2N_eV_G}{W})s_i\Big]\right. \\
\left.\left.-\alpha\Lambda z_i(1-z_i)\exp\Big[(\lambda_{m_{i+1}}-\lambda_{m_i}-\frac{4N_eV_G}{W})s_i\Big]\right)\right] \\
dz_1\ldots dz_k ds_1\ldots ds_k.
\end{aligned}
\tag{A31}
$$

Assuming again Fubini's theorem is satisfied, we move the integrals inside the series of sums, and surround the $k$-fold product. And thus, we've reduced the path integral to a $2k$- fold integral of the form

$$\int_0^t \int_0^{s_k} \cdots\cdots \int_0^{s_2} \int_0^1 \underbrace{\cdots\cdots}_{k \text{ times}} \int_0^1 \Bigg[ \prod_{i=1}^{k} z_i^2 (1-z_i) \mathcal{C}^{(3/2)}_{m_{i+1}-1}(1-2z_i) \mathcal{C}^{(3/2)}_{m_i-1}(1-2z_i)$$
$$\Bigg( 2V_G \exp\Big[ (\lambda_{m_{i+1}} - \lambda_{m_i} - \frac{2N_e V_G}{W}) s_i \Big]$$
$$- \alpha\Lambda(1-z_i) \exp\Big[ (\lambda_{m_{i+1}} - \lambda_{m_i} - \frac{4N_e V_G}{W}) s_i \Big] \Bigg) \Bigg]$$
$$dz_1 \dots dz_k ds_1 \dots ds_k. \tag{A32}$$

We can display the expression inside the brackets as

$$\prod_{i=1}^{k} \sum_{j=0}^{1} (2V_G)^{1-j} (-\alpha\Lambda)^j z_i^2 (1-z_i)^{1+j} \mathcal{C}^{(3/2)}_{m_{i+1}-1}(1-2z_i) \mathcal{C}^{(3/2)}_{m_i-1}(1-2z_i)$$
$$\cdot \exp\Big[ (\lambda_{m_{i+1}} - \lambda_{m_i} - \frac{2^{1+j} N_e V_G}{W}) s_i \Big]. \tag{A33}$$

Exchanging products and sums (Equation A27), we can rewrite Equation A32 as

$$\int_0^t \int_0^{s_k} \cdots\cdots \int_0^{s_2} \int_0^1 \underbrace{\cdots\cdots}_{k \text{ times}} \int_0^1 \sum_{j_1=0}^{1} \cdots \sum_{j_k=0}^{1} \Bigg[ \prod_{i=1}^{k} (2V_G)^{1-j_i} (-\alpha\Lambda)^{j_i} z_i^2 (1-z_i)^{1+j_i}$$
$$\mathcal{C}^{(3/2)}_{m_{i+1}-1}(1-2z_i) \mathcal{C}^{(3/2)}_{m_i-1}(1-2z_i) \Bigg]$$
$$\cdot \exp\Bigg[ \sum_{i=1}^{k} (\lambda_{m_{i+1}} - \lambda_{m_i} - \frac{2^{1+j_i} N_e V_G}{W}) s_i \Bigg]$$
$$dz_1 \dots dz_k ds_1 \dots ds_k, \tag{A34}$$

noting the first bracket is a function of $z_i$ and the exponent a function of $s_i$. We exchange the integrals and sums

$$\sum_{j_1=0}^{1} \cdots \sum_{j_k=0}^{1} \Bigg[ \prod_{i=1}^{k} (2V_G)^{1-j_i} (-\alpha\Lambda)^{j_i} \int_0^1 z_i^2 (1-z_i)^{1+j_i}$$
$$\mathcal{C}^{(3/2)}_{m_{i+1}-1}(1-2z_i) \mathcal{C}^{(3/2)}_{m_i-1}(1-2z_i) dz_i \Bigg]$$
$$\int_0^t \int_0^{s_k} \cdots\cdots \int_0^{s_2} \exp\Bigg[ \sum_{i=1}^{k} (\lambda_{m_{i+1}} - \lambda_{m_i} - \frac{2^{1+j_i} N_e V_G}{W}) s_i \Bigg] ds_1 \dots ds_k, \tag{A35}$$

and, thus, successfully disentangle the integrals over space and time.

##### A.5 Space Integrals

Focusing first on the space integral in the first set of brackets. First substitute $u_i = 1 - 2z_i$

$$
\begin{aligned}
&\int_0^1 z_i^2(1-z_i)^{1+j_i}\mathcal{C}^{(3/2)}_{m_{i+1}-1}(1-2z_i)\mathcal{C}^{(3/2)}_{m_i-1}(1-2z_i)dz_i \\
&= -\frac{1}{2^{4+j_i}}\int_1^{-1}(1-u_i)^2(1+u_i)^{1+j_i}\mathcal{C}^{(3/2)}_{m_{i+1}-1}(u_i)\mathcal{C}^{(3/2)}_{m_i-1}(u_i)du_i \\
&= \frac{1}{2^{4+j_i}}\int_{-1}^1(1-u_i^2)(1-u_i^{1+j_i})\mathcal{C}^{(3/2)}_{m_{i+1}-1}(u_i)\mathcal{C}^{(3/2)}_{m_i-1}(u_i)du_i \\
&= \frac{1}{2^{4+j_i}}\left(\int_{-1}^1(1-u_i^2)\mathcal{C}^{(3/2)}_{m_{i+1}-1}(u_i)\mathcal{C}^{(3/2)}_{m_i-1}(u_i)du_i\right. \\
&\qquad\left. - \int_{-1}^1 u_i^{1+j_i}(1-u_i^2)\mathcal{C}^{(3/2)}_{m_{i+1}-1}(u_i)\mathcal{C}^{(3/2)}_{m_i-1}(u_i)du_i\right). \qquad \text{(A36)}
\end{aligned}
$$

The first integral is the orthogonality condition of the GegenBauers,

$$
\begin{aligned}
\int_{-1}^1(1-u_i^2)\mathcal{C}^{(3/2)}_{m_{i+1}-1}(u_i)\mathcal{C}^{(3/2)}_{m_i-1}(u_i)du_i &= \delta_{m_{i+1}-1,m_i-1}\frac{2(m_{i+1}-1+1)(m_{i+1}-1+2)}{3+2(m_{i+1}-1)} \\
&= \delta_{m_{i+1},m_i}\frac{2m_i(m_i+1)}{2m_i+1}. \qquad \text{(A37)}
\end{aligned}
$$

The second integral can be made to look like the orthogonality condition using the recurrence relation of the Gegenbauers,

$$
\begin{aligned}
u_i\mathcal{C}^{(3/2)}_{m_i-1}(u_i) &= \frac{1}{3+2(m_i-1)}\left((m_i-1+1)\mathcal{C}^{(3/2)}_{m_i-1+1}(u_i)+(m_i-1+2)\mathcal{C}^{(3/2)}_{m_i-1-1}(u_i)\right) \\
&= \frac{1}{1+2m_i}\left(m_i\mathcal{C}^{(3/2)}_{m_i}(u_i)+(m_i+1)\mathcal{C}^{(3/2)}_{m_i-2}(u_i)\right), \qquad \text{(A38)}
\end{aligned}
$$

the same is true for $m_{i+1}$. With a bit of algebra, the space integral A36 is equal to

$$
\begin{aligned}
\underset{\substack{|m_{i+1}-m_i| \\ \leq 1+j_i}}{\Delta} &= \int_0^1 z_i^2(1-z_i)^{1+j_i}\mathcal{C}^{(3/2)}_{m_{i+1}-1}(1-2z_i)\mathcal{C}^{(3/2)}_{m_i-1}(1-2z_i)dz_i \\
&= \frac{1}{2^{4+j_i}}\Bigg(\delta_{m_{i+1},m_i}\frac{2m_i(m_i+1)}{2m_i+1} \\
&\qquad -\Bigg[\frac{1}{2m_i+1}\Bigg(\delta_{m_{i+1}-1,m_i}\frac{2m_i(m_i+1)(m_i+2)}{2m_i+3} \\
&\qquad\qquad +\delta_{m_{i+1},m_i-1}\frac{2(m_i-1)m_i(m_i+1)}{2m_i-1}\Bigg)\Bigg]^{1-j_i} \\
&\qquad \cdot\Bigg[\frac{1}{(2m_{i+1}+1)(2m_i+1)}\Bigg(\delta_{m_{i+1},m_i}\frac{2m_i^2(m_i+1)(m_i+2)}{2m_i+3} \\
&\qquad\qquad +\delta_{m_{i+1},m_i-2}\frac{2(m_i-1)m_i^2(m_i+1)}{2m_i-1} \\
&\qquad\qquad +\delta_{m_{i+1}-2,m_i}\frac{2m_i(m_i+1)^2(m_i+2)}{2m_i+3} \\
&\qquad\qquad +\ \delta_{m_{i+1},m_i}\frac{2(m_i-1)m_i(m_i+1)^2}{2m_i+3}\Bigg)\Bigg]^{j_i}\Bigg).
\end{aligned}
\tag{A39}
$$

Which is equal to 0 unless one of the following are true:

$$
\begin{aligned}
m_{i+1} &= m_i, \\
m_{i+1} &= m_i - 1 - j_i, \\
m_{i+1} &= m_i + 1 + j_i,
\end{aligned}
$$

for $j_i = 0$ or 1.

#### A.6 Time Integrals

Turning now to the series of time integrals on the final line of Equation A35. This, too, can be solved exactly, although a form for a general $k$ is elusive. We show the $k = 2$ case for illustration, but in practice we let Mathematica make quick work of this.

$$\begin{aligned}\int_0^t \int_0^{s_2} \exp\left[\sum_{i=1}^{2} \chi_i s_i\right] ds_1 ds_2 &= \int_0^t \int_0^{s_2} \prod_{i=1}^{2} \exp\left[\chi_i s_i\right] ds_1 ds_2 \\ &= \int_0^t \exp\left[\chi_2 s_2\right] \int_0^{s_2} \exp\left[\chi_1 s_1\right] ds_1 ds_2 \\ &= \frac{1}{\chi_1} \int_0^t \exp\left[\chi_2 s_2\right] \left(\exp\left[\chi_1 s_2\right] - 1\right) ds_2 \\ &= \frac{1}{\chi_1} \int_0^t \left(\exp\left[(\chi_1 + \chi_2) s_2\right] - \exp\left[\chi_2 s_2\right]\right) ds_2 \\ &= \frac{1}{\chi_1(\chi_1 + \chi_2)} \left(\exp\left[(\chi_1 + \chi_2) t\right] - 1\right) \\ &\qquad - \frac{1}{\chi_1 \chi_2} \left(\exp\left[\chi_2 t\right] - 1\right), \end{aligned} \tag{A40}$$

where

$$\chi_i = \lambda_{m_{i+1}} - \lambda_{m_i} - \frac{2^{1+j_i} N_e V_G}{W}. \tag{A41}$$

Attempts can be made to find common denominators and combine terms where possible, but we've not been able to find a satisfactory general solution. The number of terms grows with $\leq 2^k$ (each integral splits a term into 2, but some may combine and a lot cancel). However, none of the individual integrals are any more complicated than

$$\int_0^{s_{i+1}} \exp[\chi_i s_i] ds_i = \chi_i^{-1}(\exp[\chi_i s_{i+1}] - 1). \tag{A42}$$

The example shown assumes none of the $\chi_i$, or their sums, equal 0. If any are 0, some of the exponential terms in the intermediate steps reduce to 1, and the solution is distinct from the more general case shown (Equation A40). In fact, plugging any $\chi_i$ or $\sum_i \chi_i = 0$ into the general case, results in division by zero errors. We handled each edge case individually, for each set of $\{\chi_i\}$ that sum to zero.

##### A.7 Final Expression

Thus, we have

$$P_\alpha(y,t|x,0) = \exp\left[\frac{2N_e\alpha\Lambda}{W}\left(y\cdot\exp\left[-\frac{2N_eV_G}{W}t\right]-x\right)\right]\cdot\sum_{k=0}^{\infty}\left(\frac{8N_e^2\alpha\Lambda}{W^2}\right)^k x(1-x)$$
$$\sum_{m_1=1}^{\infty}\cdots\sum_{m_{k+1}=1}^{\infty}\mathcal{C}_{m_1-1}^{(3/2)}(1-2x)\mathcal{C}_{m_{k+1}-1}^{(3/2)}(1-2y)\exp\left[-\lambda_{m_{k+1}}t\right]$$
$$\left[\prod_{i=1}^{k+1}4\frac{2m_i+1}{m_i(m_i+1)}\right]\sum_{j_1=0}^{1}\cdots\sum_{j_k=0}^{1}\left[\prod_{i=1}^{k}(2V_G)^{1-j_i}(-\alpha\Lambda)^{j_i}\underset{\substack{|m_{i+1}-m_i|\\ \leq 1+j_i}}{\Delta}\right]$$
$$\int_0^t\int_0^{s_k}\cdots\cdots\int_0^{s_2}\exp\left[\sum_{i=1}^{k}(\lambda_{m_{i+1}}-\lambda_{m_i}-\frac{2^{1+j_i}N_eV_G}{W})s_i\right]ds_1\ldots ds_k, \tag{A43}$$

where, again, the series of integrals in the final line is solvable, but not in general. The $k$ fold product will be equal to 0 unless $|m_{i+1}-m_i|\leq 1+j_i$ **for all** $i\in 1:k$, which greatly reduces the number of terms in the series of sums which need be evaluated.

Following similar steps as above with $S_\alpha(z) = -\frac{N_e\alpha^2}{W}(1-2z)$ (under-dominant selection) we arrive at a similar expression

$$\exp\left[-\frac{N_e\alpha^2}{W}\left(y(1-y)-x(1-x)\right)\right]\sum_{k=0}^{\infty}\left(-\frac{N_e\alpha^2}{W}\right)^k x(1-x)\sum_{m_1=1}^{\infty}\cdots\sum_{m_{k+1}=1}^{\infty}$$
$$\mathcal{C}_{m_1-1}^{(3/2)}(1-2x)\mathcal{C}_{m_{k+1}-1}^{(3/2)}(1-2y)\exp\left[-\lambda_{m_{k+1}}t\right]\left[\prod_{i=1}^{k+1}4\frac{2m_i+1}{m_i(m_i+1)}\right]$$
$$\sum_{j_1=0}^{1}\cdots\sum_{j_k=0}^{1}\left[\prod_{i=1}^{k}\left(\frac{N_e\alpha^2}{2W}\right)^{j_i}\int_0^1 z_i^2(1-z_i)^2(1-2z_i)^{2j_i}\right.$$
$$\left.\mathcal{C}_{m_{i+1}-1}^{(3/2)}(1-2z_i)\mathcal{C}_{m_i-1}^{(3/2)}(1-2z_i)dz_i\right]$$
$$\int_0^t\int_0^{s_k}\cdots\cdots\int_0^{s_2}\exp\left[\sum_{i=1}^{k}(\lambda_{m_{i+1}}-\lambda_{m_i})s_i\right]ds_1\ldots ds_k. \tag{A44}$$

Noting that the integrals on the final of sums has been moved outside of the sums over the $j_i$s (because $\chi_i$ is no longer a function of $j_i$).

### B Path Integral Expression of the Full Model

After rescaling time to genomic units of $2N_e$ generations, the full selection model is

$$\begin{aligned} S_\alpha(z,s) &= 2N_e\alpha\frac{\Lambda}{W}\exp\left[-\frac{2N_eV_G}{W}s\right] - \frac{N_e\alpha^2}{W}\left(1-\frac{\Lambda^2}{W}\exp\left[-\frac{4N_eV_G}{W}\right]\right)(1-2z) \\ &= 2N_e\alpha\frac{\Lambda}{W}\exp\left[-\frac{2N_eV_G}{W}s\right] - \frac{N_e\alpha^2}{W}(1-2z) + \frac{N_e\alpha^2\Lambda^2}{W^2}\exp\left[-\frac{4N_eV_G}{W}\right](1-2z). \end{aligned} \tag{A45}$$

And so, the Ito Integral in Equation A8 is

$$\begin{aligned} \int_0^t S_\alpha(z_s,s)dz_s &= \int_0^t 2N_e\alpha\frac{\Lambda}{W}\exp\left[-\frac{2N_eV_G}{W}s\right] - \frac{N_e\alpha^2}{W}(1-2z) \\ &\qquad + \frac{N_e\alpha^2\Lambda^2}{W^2}\exp\left[-\frac{4N_eV_G}{W}\right](1-2z)dz_s \\ &= \int_0^t 2N_e\alpha\frac{\Lambda}{W}\exp\left[-\frac{2N_eV_G}{W}s\right]dz_s + \int_0^t -\frac{N_e\alpha^2}{W}(1-2z)dz_s \\ &\qquad + \int_0^t \frac{N_e\alpha^2\Lambda^2}{W^2}\exp\left[-\frac{4N_eV_G}{W}s\right](1-2z)dz_s. \end{aligned} \tag{A46}$$

The first integral is the same we saw in the simplified model above

$$\begin{aligned} &\int_0^t 2N_e\alpha\frac{\Lambda}{W}\exp\left[-\frac{2N_eV_G}{W}s\right]dz_s \\ &\qquad = \frac{2N_e\alpha\Lambda}{W}\left(y\cdot\exp\left[-\frac{2N_eV_G}{W}t\right] - x + \frac{2N_eV_G}{W}\int_0^t z_s\exp\left[-\frac{2N_eV_G}{W}s\right]ds\right), \end{aligned} \tag{A47}$$

the second is the same as in the constant stabilizing selection case (not shown):

$$\int_0^t -\frac{N_e\alpha^2}{W}(1-2z)dz_s = -\frac{N_e\alpha^2}{W}\left(y(1-y) - x(1-x) + \int_0^t z_s(1-z_s)ds\right), \tag{A48}$$

and the third

$$\begin{aligned} \int_0^t \frac{N_e\alpha^2\Lambda^2}{W^2}\exp\left[-\frac{4N_eV_G}{W}\right](1-2z)dz_s &= \frac{N_e\alpha^2\Lambda^2}{W^2}\left(y(1-y)\exp\left[-\frac{4N_eV_G}{W}t\right] - x(1-x)\right. \\ &\qquad \left. + \left(\frac{4N_eV_G}{W}+1\right)\int_0^t z_s(1-z_s)\exp\left[-\frac{4N_eV_G}{W}s\right]ds\right). \end{aligned} \tag{A49}$$

So the ratio inside of the path integral (Equation A8) is

$$\begin{aligned}
\rho_{\alpha,0} =& \exp\left[\frac{2N_e\alpha\Lambda}{W}\left(y\cdot\exp\left[-\frac{2N_eV_G}{W}t\right]-x\right)\right.\\
&\qquad -\frac{N_e\alpha^2}{W}\left(y(1-y)-x(1-x)\right)\\
&\qquad \left.+\frac{N_e\alpha^2\Lambda^2}{W^2}\left(y(1-y)\exp\left[-\frac{4N_eV_G}{W}t\right]-x(1-x)\right)\right]\\
&\cdot\exp\left[\int_0^t\frac{2N_eV_G}{W}\exp\left[-\frac{2N_eV_G}{W}s\right]z_s\right.\\
&\qquad -\frac{N_e\alpha^2}{W}z_s(1-z_s)\\
&\qquad +\left(\frac{4N_eV_G}{W}+1\right)\exp\left[-\frac{4N_eV_G}{W}s\right]z_s(1-z_s)\\
&\qquad -\frac{N_e^2\alpha^4}{2W^2}z_s(1-z_s)(1-2z_s)\\
&\qquad +\frac{2N_e^2\alpha^3\Lambda}{W^2}\exp\left[-\frac{2N_eV_G}{W}s\right]z_s(1-z_s)(1-2z_s)\\
&\qquad -\frac{2N_e^2\alpha^2\Lambda^2}{W^2}\exp\left[-\frac{4N_eV_G}{W}s\right]z_s(1-z_s)\\
&\qquad +\frac{N_e^2\alpha^4\Lambda^2}{W^3}\exp\left[-\frac{4N_eV_G}{W}s\right]z_s(1-z_s)(1-2z_s)^2\\
&\qquad -\frac{2N_e^2\alpha^3\Lambda^3}{W^3}\exp\left[-\frac{6N_eV_G}{W}s\right]z_s(1-z_s)(1-2z_s)\\
&\qquad \left.-\frac{N_e^2\alpha^4\Lambda^4}{2W^4}\exp\left[-\frac{8N_eV_G}{W}s\right]z_s(1-z_s)(1-2z_s)ds\right]\\
=& \exp[\ldots]\cdot\exp\left[\int_0^t\sum_{j=1}^{9}\phi_j(z_s,s)ds\right]. \qquad \text{(A50)}
\end{aligned}$$

Each of these terms can be solved using the same methods as above. However, because each of the terms in $\phi_j$ are rather unique, we are unable to neatly combine them into a single function of $j$ (as we did in Equation A33). In the full solution, the series of sums over $j_i$ range from 1 to 9, increasing the number of terms to be evaluated by a factor of $4.5^k$. For this reason, we chose to focus on the simplified model, which is already computationally intensive as it is. For a single combination of $N_e$, $V_G$ and $W$ parameters, it took over 5 hours just to compute the sums in Equation A43 (running on 12 kernels in parallel). Furthermore, an additional 5 hours was required to generate the possible edge cases of the time integrals, and 1 hour to pre-compute the time integrals for each case, but this does not need to be repeated for different parameter combinations.

### C Expected FDR

The expected false discovery rate is the probability that an allele meeting or exceeding the AFC threshold ($y \geq \Psi_x$) does not contribute to the trait under selection ($\alpha = 0$). We can write an expression for this probability as a ratio of Bayes' theorem,

$$\frac{P(\alpha = 0|y \geq \Psi_x)}{P(\alpha \neq 0|y \geq \Psi_x)} = \frac{P(y \geq \Psi_x|\alpha = 0)P(\alpha = 0)}{P(y \geq \Psi_x|\alpha \neq 0)P(\alpha \neq 0)}. \tag{A51}$$

Equations of this form are widely used in clinical settings (Habibzadeh and Habibzadeh, 2019), and the fraction

$$\frac{P(y \geq \Psi_x|\alpha = 0)}{P(y \geq \Psi_x|\alpha \neq 0)} \tag{A52}$$

is known as the positive likelihood ratio.

Recognizing that

$$P(\alpha \neq 0|y \geq \Psi_x) = 1 - P(\alpha = 0|y \geq \Psi_x), \tag{A53}$$

this can be rewritten as

$$P(\alpha = 0|y \geq \Psi_x) = \left[1 + \frac{P(y \geq \Psi_x|\alpha \neq 0)P(\alpha \neq 0)}{P(y \geq \Psi_x|\alpha = 0)P(\alpha = 0)}\right]^{-1}. \tag{A54}$$

We focus on a given starting frequency $x$, and assume that $U_\alpha$ mutations appear per haplotype per generation with effect $\alpha$. We show later how to extend these results to a genome wide scan and an arbitrary distribution of effect sizes. With our simplifying assumptions, we can rewrite Equation A54 as

$$P(\alpha = 0|y \geq \Psi_x, x) = \left[1 + \frac{P(y \geq \Psi_x|\alpha, x)P(\alpha|x)}{P(y \geq \Psi_x|0, x)P(0|x)}\right]^{-1} \tag{A55}$$

By construction (Section 4.3),

$$\begin{aligned} P(y \geq \Psi_x|\alpha, x) &= Q_{x,\alpha}, \\ P(y \geq \Psi_x|0, x) &= q. \end{aligned} \tag{A56}$$

Assuming the starting population is in equilibrium, we can find expressions for the probabilities that an allele with starting frequency $x$ has a certain effect size using theory of random sums. For example, the expected number of alleles segregating at frequency $x$ with effect size $\alpha$ can be expressed as

$$E[\#_{\alpha|x}] = E[\sum^{n_x} \mathbf{1}_\alpha], \tag{A57}$$

where $n_x$ is the total number of alleles segregating at frequency $x$ and $\mathbf{1}_\alpha$ is a Bernoulli random variable that takes value 1 with probability $P(\alpha|x)$. Assuming independence between sites, these are independent. And for a fixed $x$ and $\alpha$, these are identically distributed. Then the expectation of the random sum can be found using Wald's Identity

$$E[\sum^{n_x} \mathbf{1}_\alpha] = E[n_x]P(\alpha|x) \tag{A58}$$

(Resnick, 2013, Chapter 1.8.1). Then,

$$P(\alpha|x) = \frac{E[n_{\alpha|x}]}{E[n_x]}. \tag{A59}$$

The expected number of neutral alleles segregating at frequency between $x$ and $x + dx$ is $2NU_0\Phi_0(x)dx$, where $U_0$ is the number of neutral mutations per haplotype per generation and $\Phi$ is the density of the expected amount of time a mutation spends at frequency $x$ before fixation or loss (Kimura, 1969; Simons *et al.*, 2018; Yair and Coop, 2022). Similarly, the expected number of trait-contributing alleles starting at frequency between $x$ and $x+dx$ with effect $\alpha$ is $2NU_\alpha\Phi_\alpha(x)dx$. The sojourn time can be found by integrating the transition probability from $t = 0$ to $\infty$ or by evaluating a general solution (Ewens, 1963; Kimura, 1969; Maruyama, 1974; Ewens, 2004, Chapter 4.4). Substituting into Equation A55, we have

$$P(\alpha = 0|y \geq \Psi_x, x) = \left[1 + \frac{Q_{x,\alpha}U_\alpha\Phi_\alpha(x)}{qU_0\Phi_0(x)}\right]^{-1}. \tag{A60}$$

This result can be extended to an arbitrary distribution of effect sizes for trait contributing alleles (denoted $\kappa(\alpha)$) by recognizing

$$\begin{aligned} P(y \geq \Psi_x|\alpha \neq 0, x)P(\alpha \neq 0|x) &= \int P(y \geq \Psi_x|\varepsilon, x)P(\varepsilon|x)d\varepsilon \\ &\propto \int Q_{x,\varepsilon}2NU\kappa(\varepsilon)\Phi_\varepsilon(x)d\varepsilon. \end{aligned} \tag{A61}$$

Here, $U$ is the rate at which trait contributing mutations arise (so $U_\alpha = U\kappa(\alpha)$). This is equivalent to Equation A60 when

$$\kappa(\varepsilon) = \delta(\varepsilon - \alpha), \tag{A62}$$

where $\delta(\cdot)$ is the Dirac delta function. In our simulations (Section 4.2), trait contributing mutations had effect $\pm\alpha$ with equal probability. In this case,

$$\kappa(\varepsilon) = \frac{1}{2}\delta(\varepsilon - \alpha) + \frac{1}{2}\delta(\varepsilon + \alpha), \tag{A63}$$

and

$$P(\alpha = 0|y \geq \Psi_x, x) = \left[1 + \frac{Q_{x,\alpha}U/2\Phi_\alpha(x) + Q_{x,-\alpha}U/2\Phi_{-\alpha}(x)}{qU_0\Phi_0(x)}\right]^{-1}. \quad (A64)$$

Because $\Phi$ for symmetric underdominance is a function of $\alpha^2$, $\Phi_\alpha(x) = \Phi_{-\alpha}(x)$. Thus,

$$P(\alpha = 0|y \geq \Psi_x, x) = \left[1 + \frac{Q_{x,\alpha} + Q_{x,-\alpha}}{q} \cdot \frac{U\Phi_\alpha(x)}{2U_0\Phi_0(x)}\right]^{-1}. \quad (A65)$$

Finally, given an observed distribution of starting frequencies of alleles conditioned on exceed our AFC threshold (denoted $\text{SFS}(x|y \geq \Psi_x)$), we can find the expected false discovery rate of a genome wide scan by weighting the expected false discovery rate at each $x$ by the proportion of candidate alleles at that frequency:

$$P(\alpha = 0|y \geq \Psi) = \frac{1}{\sum_x \text{SFS}(x|y \geq \Psi_x)} \sum_x P(\alpha = 0|y \geq \Psi_x, x)\text{SFS}(x|y \geq \Psi_x). \quad (A66)$$

So far, we have only considered testing the derived allele for extreme AFCs. In practice, however, researchers will restrict their analysis to alleles which increase in frequency (the ”rising allele”; e.g., Tobler *et al.*, 2014; Barghi *et al.*, 2019). This does not differentiate whether a given allele is positively selected, or its alternate allele is negatively selected (or *vice versa*). In fact, the two scenarios are equivalent

$$\begin{aligned} M_{\text{alt}} &= -M \\ V_{\text{alt}} &= V. \end{aligned} \quad (A67)$$

One can imagine a negatively selected allele starting at frequency $1 - x$ decreasing in frequency such that its alternate allele's ending frequency exceeds $\Psi_x$ and is labeled a target of selection. This scenario can be easily accounted for in the case of symmetric underdominance (so that $\Phi_\alpha(x) = \Phi_{-\alpha}(x)$) and a distribution of effect sizes symmetric about 0 ($\kappa(\alpha) = \kappa(-\alpha)$). In this case, $\Phi_\alpha(x)$ is replaced with $\Phi_\alpha(x) + \Phi_\alpha(1 - x)$ in the above equations. For example, Equation A65 becomes

$$P(\alpha = 0|y \geq \Psi_x, x, y \geq x) = \left[1 + \frac{Q_{x,\alpha} + Q_{x,-\alpha}}{q} \cdot \frac{U\left(\Phi_\alpha(x) + \Phi_\alpha(1 - x)\right)}{2U_0\left(\Phi_0(x) + \Phi_0(1 - x)\right)}\right]^{-1}. \quad (A68)$$

We can also derive a similar expression under more general conditions (namely, when $\Phi$ and $\kappa$ are not symmetric around $\alpha = 0$) by extending the probability that an allele at starting frequency $x$ has effect $\alpha$, $P(\alpha|x)$, to also include alleles at frequency $1 - x$ with effect $-\alpha$

$$\begin{aligned} P(\alpha, x \text{ or } -\alpha, 1 - x|x \text{ or } 1 - x, y \geq x) \propto\ & 2NU\kappa(\alpha)\Phi_\alpha(x)P(y \geq x|\alpha, x) \\ &+ 2NU\kappa(-\alpha)\Phi_{-\alpha}(1 - x)P(y \geq x|\alpha, x). \end{aligned} \quad (A69)$$

The probability on the left-hand side may require some explanation. This is the probability that an rising allele ($y \geq x$) at frequency $x$ has effect $\alpha$, or is the alternate of an allele at frequency $1-x$ with effect $-\alpha$. Because we are conditioning on the rising allele, we must also modify the probability of exceeding the AFC threshold

$$P(y \geq \Psi_x|\alpha, x, y \geq x) = \frac{P(y \geq \Psi_x|\alpha, x)}{P(y \geq x|\alpha, x)} = \frac{Q_{x,\alpha}}{P(y \geq x|\alpha, x)}. \tag{A70}$$

So,

$$P(y \geq \Psi_x|\alpha \neq 0, x, y \geq x)P(\alpha \neq 0|x, y \geq x) \propto \int Q_{x,\varepsilon} 2NU \left(\kappa(\varepsilon)\Phi_\varepsilon(x) + \kappa(-\varepsilon)\Phi_{-\varepsilon}(1-x)\right) d\varepsilon. \tag{A71}$$

Which is, the same result we would get if we had not conditioned on $y \geq x$ (i.e., if we had we tested every allele, regardless of whether it increased or decreased in frequency). This makes intuitive sense, because alleles that decreased in frequency would not be discovered as candidate targets of selection and, thus, would not contribute to the false discovery rate.

### D Supplemental Figures

#### D.1 Convergence

In this section we show how quickly the path integral solution converges under different strengths of selection, genetic variances, lengths of time, and values of $m_{\max}$ and $k_{\max}$.

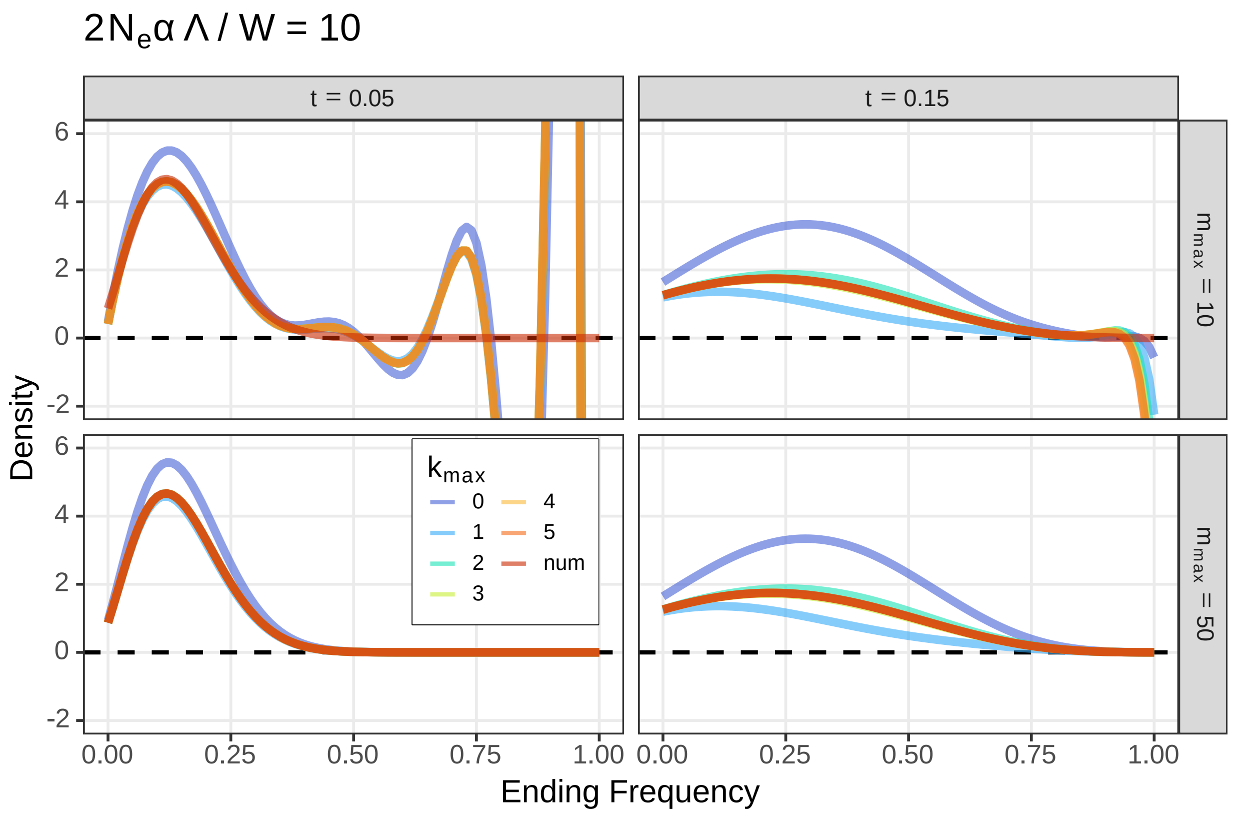


Figure S1: Greater $k_{\max}$ is needed for longer times, while greater $m_{\max}$ is needed for shorter times. We show the distribution of ending frequencies of an allele with $x = 0.1$ under strong selection ($2N_e\alpha\Lambda/W = 10$) for different amounts of time (columns). Values of $m_{\max}$ are varied along the rows and different values of $k_{\max}$ are represented by the colored lines. The numerical solution is shown as a red line. Model parameters: $\alpha = 0.01$, $N_e = 500$, $V_G = 0.001$, $W = 1$, $\Lambda = 1$

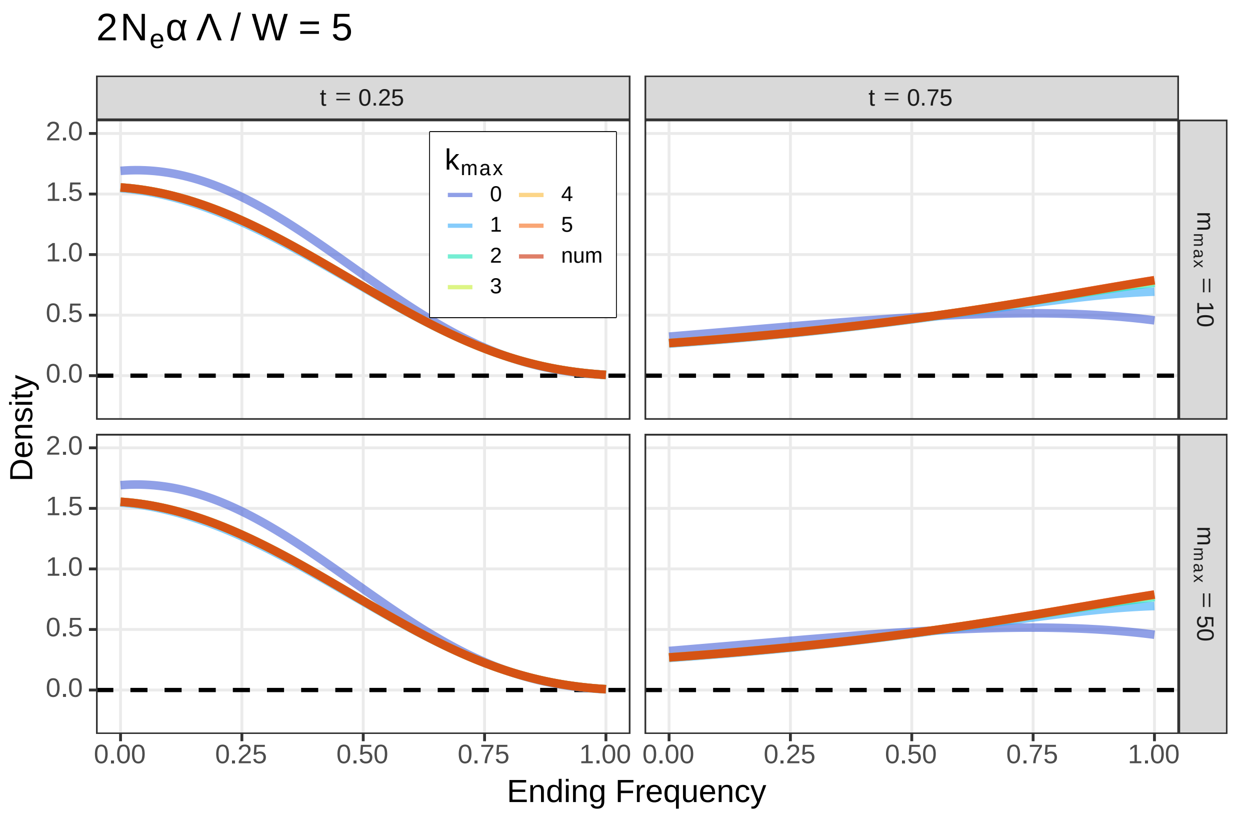


Figure S2: Smaller values of $m_{\max}$ and $k_{\max}$ are sufficient for longer times and weaker selection. We show the distribution of ending frequencies of an allele with $x = 0.1$ under weaker selection selection ($2N_e\alpha\Lambda/W = 5$, c.f. Figure S1) for different amounts of time (columns). Note the time scales are greater than in Figures S1, S3, S4. Values of $m_{\max}$ are varied along the rows and different values of $k_{\max}$ are represented by the colored lines. The numerical solution is shown as a red line. Model parameters: $\alpha = 0.005$, $N_e = 500$, $V_G = 0.001$, $W = 1$, $\Lambda = 1$.

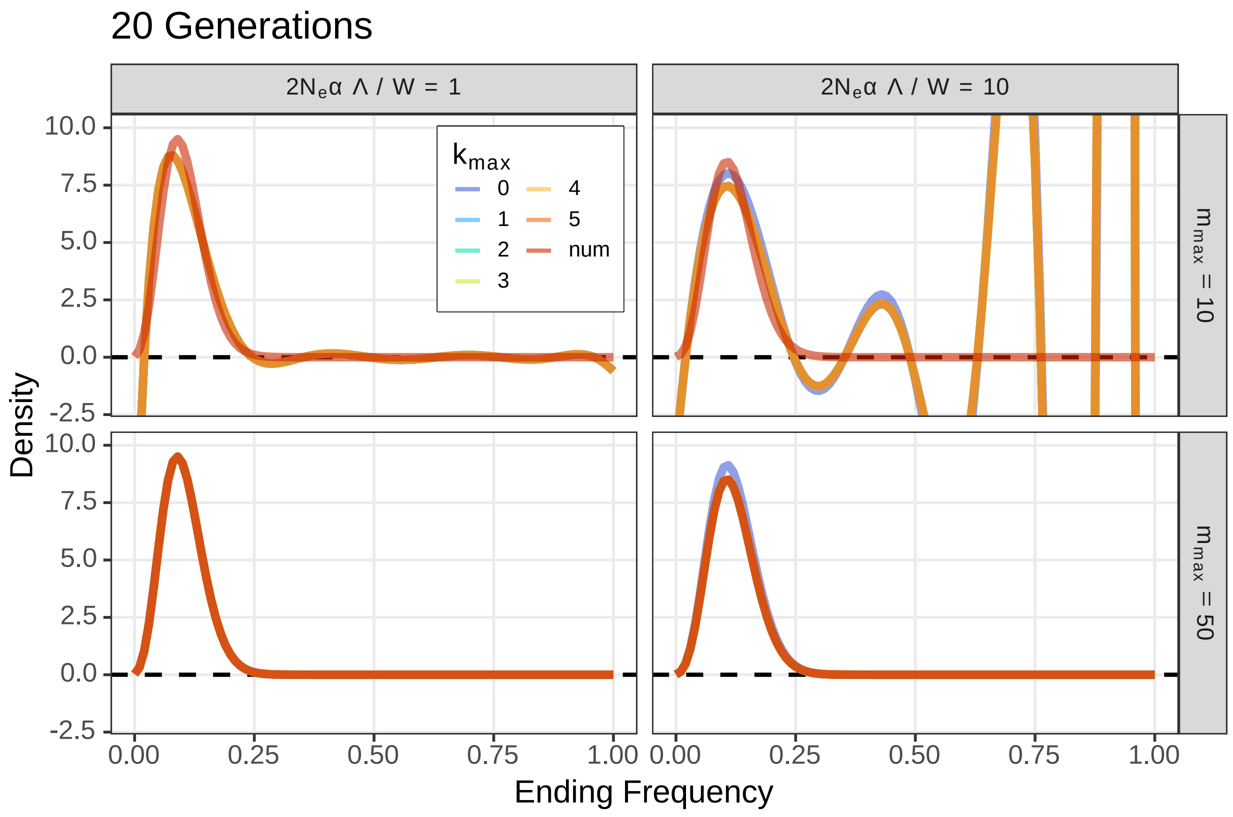


Figure S3: Convergence for small times is dominated by $m_{\max}$. We show the distribution of ending frequencies of an allele with $x = 0.1$ under weak and strong selection (columns) for 20 generations ($t = 0.02$). Values of $m_{\max}$ are varied along the rows and different values of $k_{\max}$ are represented by the colored lines. The smaller value of $m_{\max}$ is not sufficient for either strength of selection, however convergence is much worse for stronger selection. For the short time scales shown here, there is very good convergence with small $k_{\max}$ ($\approx 2$, c.f. S1). The numerical solution is shown as a red line. Model parameters: $N_e = 500$, $V_G = 0.001$, $W = 1$, $\Lambda = 1$.

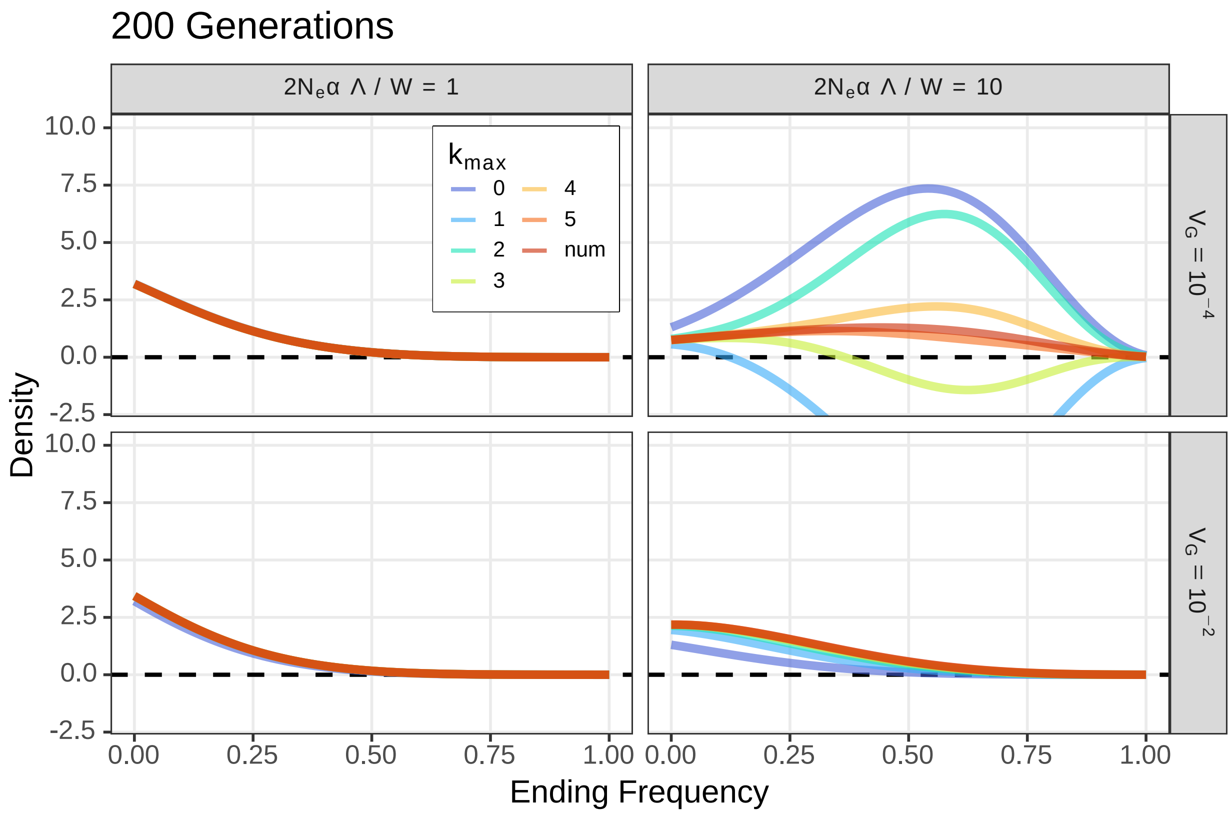


Figure S4: The rate of convergence in $k_{\max}$ is determined by both the strength of selection and its rate of decay. Greater $k_{\max}$ is needed for strong selection with small genetic variance (small rate of decay). Convergence is greatly increased for either weaker selection or greater genetic variance. We show the distribution of ending frequencies of an allele with $x = 0.1$ under various strengths of selection (columns) for 200 generations ($t = 0.2$). $V_G$ is varied along the rows and different values of $k_{\max}$ are represented by the colored lines. The numerical solution is shown as a red line. Model parameters: $N_e = 500$, $m_{\max} = 50$, $W = 1$, $\Lambda = 1$.

#### D.2 Simulation Results

In this section, we show comparisons of analytic and simulation results for a few combinations of $\alpha$ and $U$.

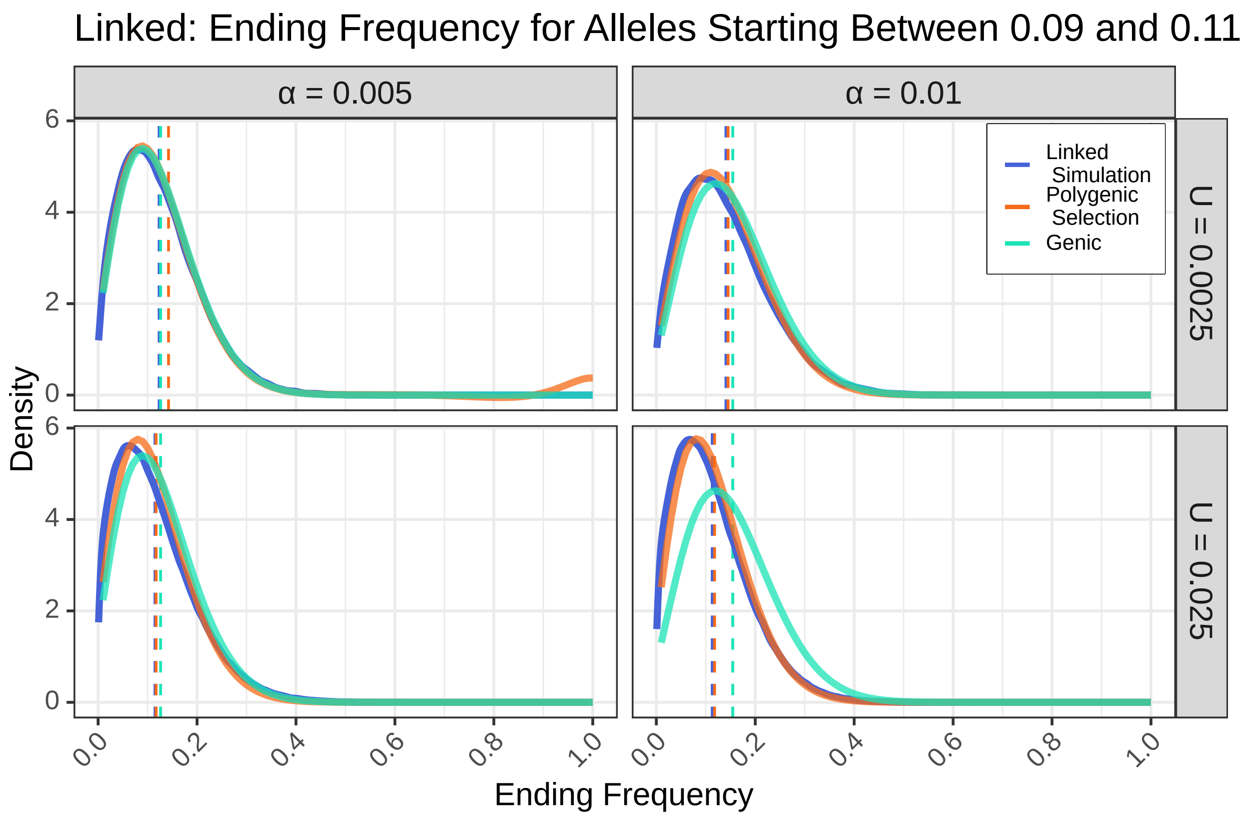


Figure S5: Our analytic solution well approximates the allele dynamics found through expensive linked simulations. For small $\alpha$ and $U$ (small $V_G$, small rate of decay in the strength of selection) the genic solution is also a good approximation. Although it is less appropriate for larger $\alpha$ and $U$ (larger $V_G$, see also Figure 2). Our solution for the time dependent selection experienced by a focal trait contributing allele after a sudden shift in optimum also performs worse under this parameter regime, but for a different reason. Our solution has slightly less heavy tails, representing AFC due to LD with either negatively selected, or more positively selected genomic backgrounds. We show the distribution of ending frequencies of a trait contributing allele starting at $x = 0.1$ after 50 generations ($t = 0.05$) of selection. Vertical dotted lines are the expectation (analytic solutions) or empirical mean (simulation) ending allele frequency. Model parameters: $N = N_e = 500$, $W = 1$, $\Lambda = 1$, $k_{\max} = 5$, $m_{\max} = 50$, $V_G \approx 0.0022, 0.0071, 0.057, 0.020$ moving clockwise starting from the top left.

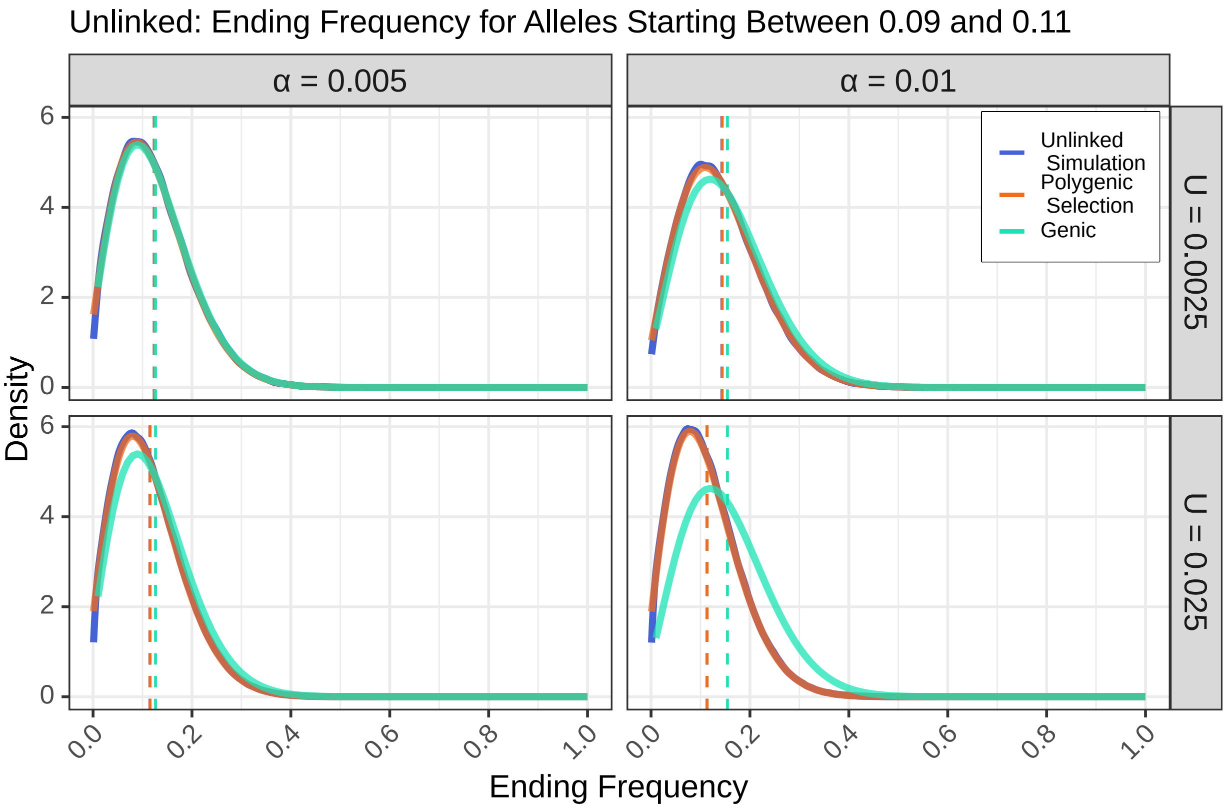


Figure S6: Our analytic solution well approximates the allele dynamics found through unlinked simulations even better than linked simulations (c.f. Figure S5). For small $\alpha$ and $U$ (small $V_G$, small rate of decay in the strength of selection) the genic solution is, again, a good approximation, but not for larger $\alpha$ and $U$ (larger $V_G$, see also Figure 2). In the absence of linkage and LD, we see good agreement between our solution and simulations for every parameter regime tested. We show the distribution of ending frequencies of a trait contributing allele starting at $x = 0.1$ after 50 generations ($t = 0.05$) of selection. Vertical dotted lines are the expectation (analytic solutions) or empirical mean (simulation) ending allele frequency. Model parameters: $N = N_e = 500$, $W = 1$, $\Lambda = 1$, $k_{\max} = 5$, $m_{\max} = 50$, $V_G \approx 0.0023, 0.0074, 0.074, 0.023$ moving clockwise starting from the top left..

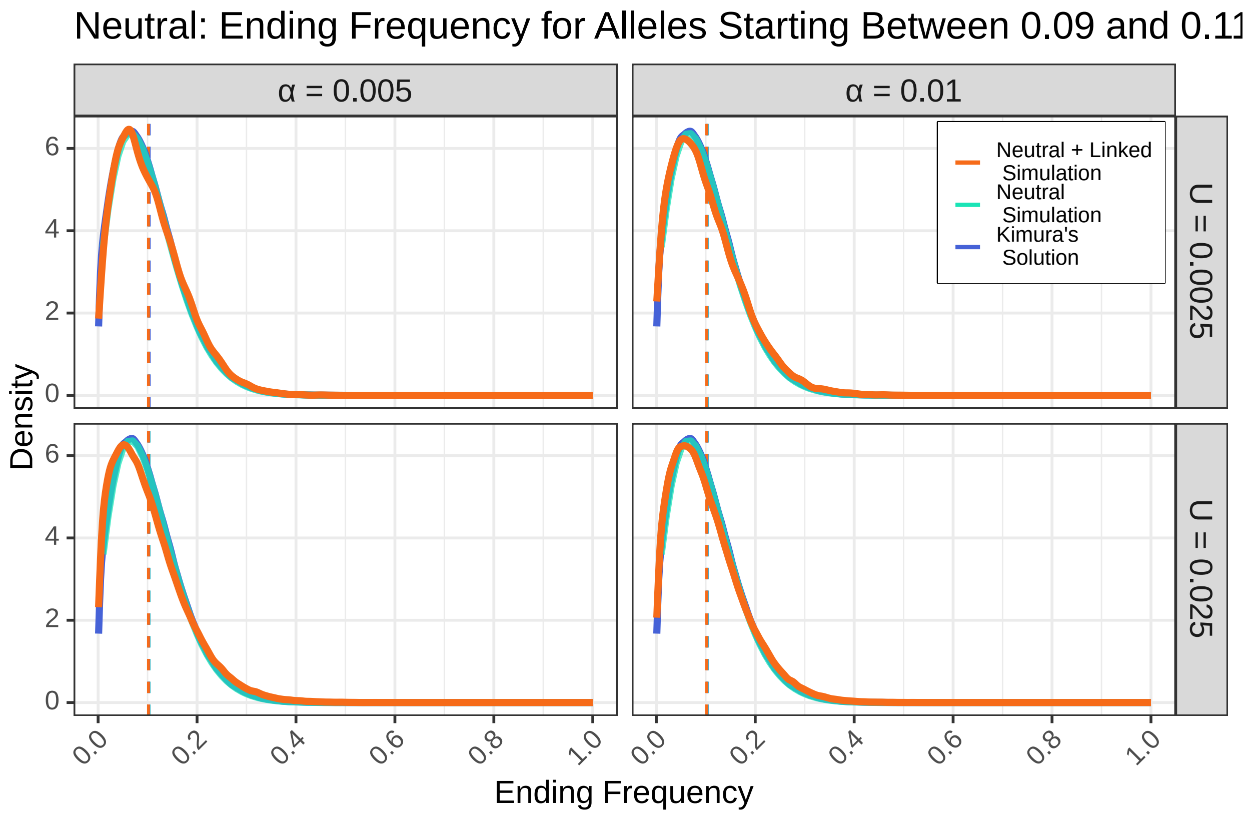


Figure S7: Kimura's neutral solution (Equation 2) well approximates the neutral allele dynamics found through both linked and unlinked simulations. The effects of linked selection are greatest for high $\alpha$ and $U$. The linked simulation distribution has slightly less heavy tails, representing AFC due to linked selection. We show the distribution of ending frequencies of a neutral allele starting at $x = 0.1$ after 50 generations ($t = 0.05$) of selection. Vertical dotted lines are the expectation (analytic solutions) or empirical mean (simulation) ending allele frequency. Model parameters: $N = N_e = 500$, $W = 1$, $\Lambda = 1$, $k_{\max} = 5$, $m_{\max} = 50$.

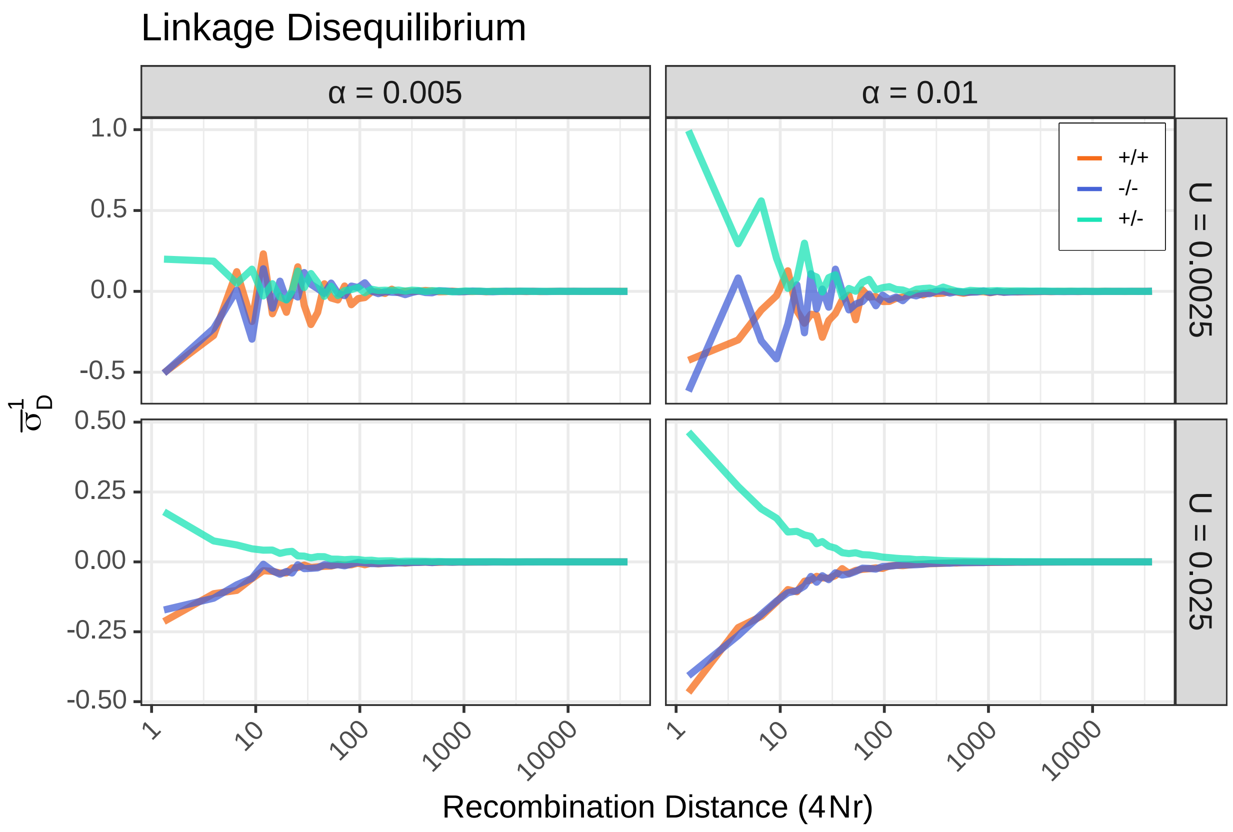


Figure S8: We observe linkage disequilibrium among trait contributing alleles (Bulmer, 1971) in our linked simulations. This violates the assumptions of our selection model, and causes the distribution of ending frequencies found through linked simulation to be heavier tailed than predicted (see Figures 3, S5, S7) We show the average linkage disequilibrium ($\sigma_D^1 = \sum_{ij} D_{ij} / \sum_{ij} \pi_{2_{ij}}$ among trait contributing alleles with positive effect sizes (+/+), negative effect sizes (−/−) and between alleles of opposite effects (+/−) as a function of recombination distance measured in the population scaled recombination rate ($\rho = 4Nr$). When $U$ is small, there may be few observations for a given recombination distance, leading to the noise seen. Model parameters: $N = 500$, $W = 1$, $\Lambda = 1$

#### D.3 Figure 4 Error

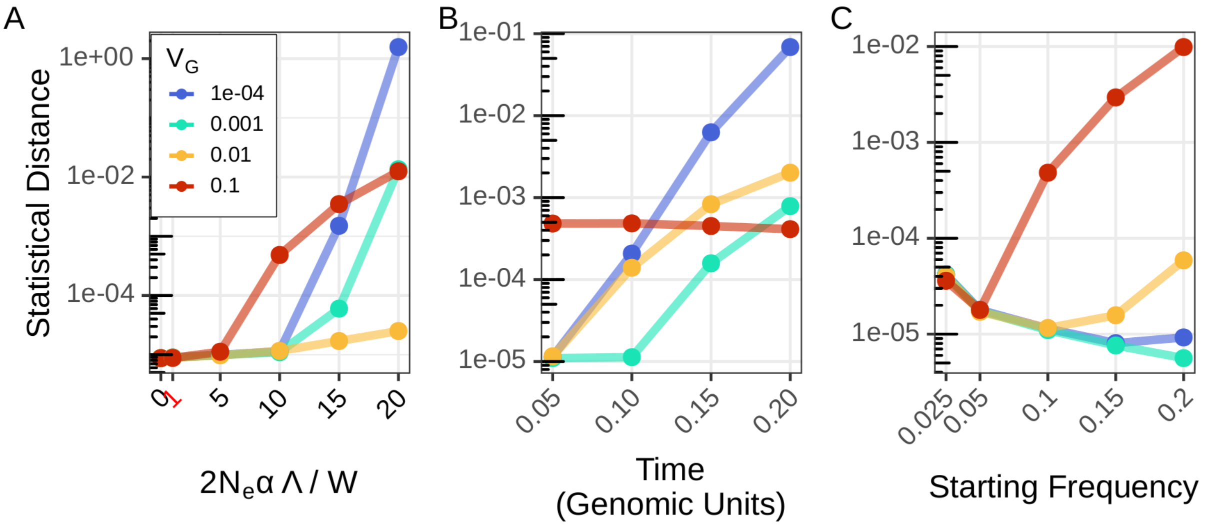


Figure S9: Error remains low across a broad range of parameter space. Here we show the statistical distance between path integral solution and numerical solution for each of the data points in Figure 4. $2N_e\alpha\Lambda/W = 10$, $t = 0.05$, $x = 0.1$ unless otherwise stated. The initial strength of selection (A), the time elapsed (B), and the starting frequency (C) are varied along the x-axis. Colors denote different values of $V_G$. Error only exceeds $10^{-2}$ when $2N_e\alpha\Lambda/W = 20$ or when $V_G$ is small and $t$ is large. Error is largest for small $V_G$, except when $x$ is large.

#### D.4 Alternative Parameter Regime Error

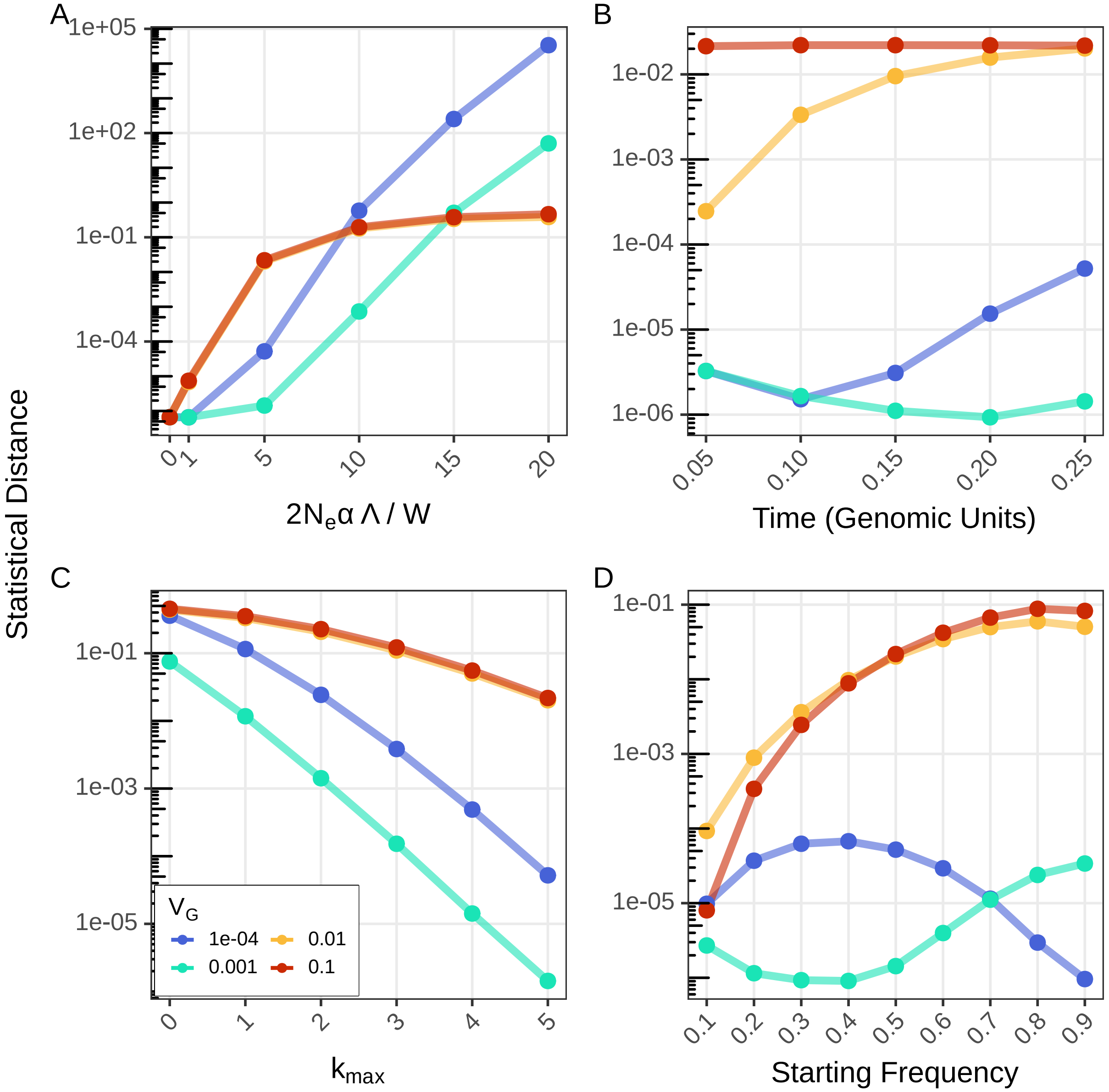


Figure S10: Error is largest for strong selection, and small $V_G$. Here we show error under a different parameter regime than elsewhere explored ($x = 0.5$, $2N_e\alpha\Lambda/W = 5$, $t = 0.25$ unless varied). Note that these are generally higher starting frequencies, longer times, and weaker selection than our main results. The initial strength of selection (A), time elapsed (B), value of $k_{\max}$ (C) and starting frequency (D) is varied along the x-axis. Model parameters: $\alpha = 0.005$, $N_e = 500$, $W = 1$, $\Lambda = 1$, $m_{\max} = 50$, $k_{\max} = 5$.

#### D.5 RFS

In this section we show the P(detected) numbers used to calculate the RFS plots (Figures 4 and 5) and their associated errors. As well as more results concerning the RFS.

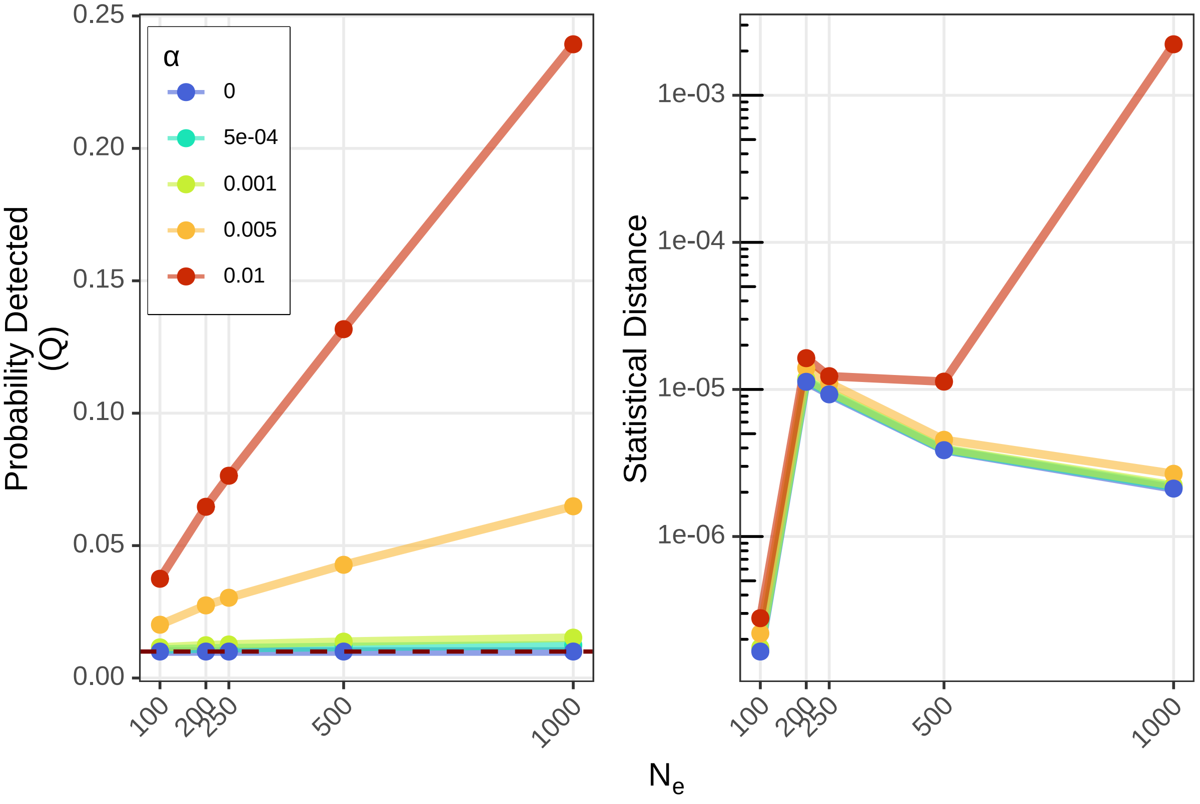


Figure S11: Probability that a trait contributing allele exceeds a 99% neutral AFC threshold after 50 generations with starting frequency $x = 0.1$. The effect size, $\alpha$, is represented by the different colors, and the effective population size, $N_e$, is varied along the x-axis. The probability is greatest when $N_e$ and $\alpha$ are large. The horizontal dotted line is the probability that a neutral allele could exceed the AFC threshold. Error associated with each of the parameter combinations shown on the left. Model parameters: $x = 0.1$, $W = 1$, $\Lambda = 1$, $V_G = 0.001$, $k_{\max} = 5$, $m_{\max} = 50$.

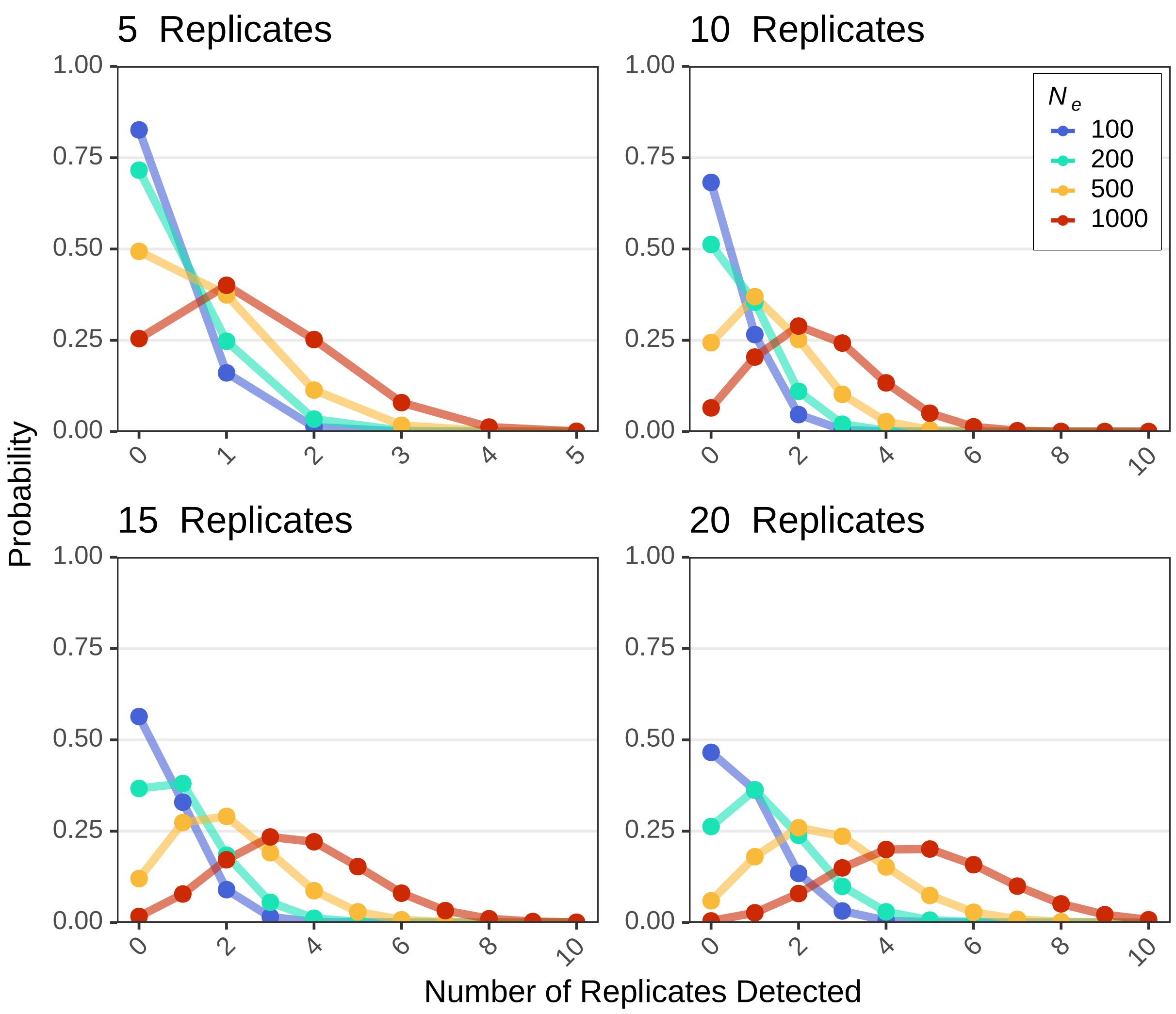


Figure S12: Both the number and size of replicate populations have an effect on the expected RFS of alleles with $\alpha = 0.01$ that exceed a 99% neutral AFC threshold after 50 generations of selection at least once among the replicate populations. The probability that an allele is detected in at least one replicate is shown in the legend. Number of replicates is varied over the different facets. Colors denote the different values of $N_e$. $2N_e\alpha\Lambda/W$ ranges from 2 to 20. Model parameters: $\alpha = 0.01$, $W = 1$, $\Lambda = 1$, $V_G = 0.001$, $k_{\max} = 5$, $m_{\max} = 50$.
